# Light-activated tetrazines enable live-cell spatiotemporal control of bioorthogonal reactions

**DOI:** 10.1101/2020.12.01.405423

**Authors:** Luping Liu, Dongyang Zhang, Mai Johnson, Neal K. Devaraj

## Abstract

Bioorthogonal ligations encompass coupling chemistries that have considerable utility in living systems.^1–3^ Among the numerous bioorthogonal chemistries described to date, cycloaddition reactions between tetrazines and strained dienophiles are widely used in proteome, lipid, and glycan labeling due to their extremely rapid kinetics.^4,5^ In addition, a variety of functional groups can be released after the cycloaddition reaction,^6,7^ and drug delivery triggered by *in vivo* tetrazine ligation^8^ is in human phase I clinical trials.^9^ While applications of tetrazine ligations are growing in academia and industry, it has so far not been possible to control this chemistry to achieve the high degrees of spatial and temporal precision necessary for modifying mammalian cells with single-cell resolution. Here we demonstrate visible light-activated formation of tetrazines from photocaged dihydrotetrazines, which enables live-cell spatiotemporal control of rapid biorthogonal cycloaddition reactions between tetrazines and dienophiles such as *trans-cyclooctenes* (TCOs). Photocaged dihydrotetrazines are stable in conditions that normally degrade tetrazines, enabling efficient early-stage incorporation of bioorthogonal handles into biomolecules such as peptides. Photocaged dihydrotetrazines allow the use of non-toxic visible light to trigger tetrazine ligations on live mammalian cells. By tagging reactive phospholipids with fluorophores, we demonstrate modification of HeLa cell membranes with single-cell spatial resolution. Finally, we show that photo-triggered therapy is possible by coupling tetrazine photoactivation with strategies that uncage prodrugs in response to tetrazine ligation, opening up new methods for photopharmacology and precision drug delivery using bioorthogonal chemistry.

---

Bioorthogonal inverse electron demand Diels-Alder reactions between tetrazines and dienophiles have found widespread use in chemical biology and material science, since their introduction in 2008.^10,11^ For example, tetrazine ligations have been used in whole animal proteome labeling,^12^ the capture of circulating tumor cells,^13^ tracking lipid modifications,^14^ and imaging glycosylation.^15^ “Click to release” strategies, which use dienophiles capable of releasing functional groups upon cycloaddition with tetrazine, have been exploited for tumor imaging,^16^ drug delivery,^17^ and controlling enzyme activity.^18^ As applications rapidly expand, there is a growing need for methods that can precisely control the reaction in the presence of live cells. Due to its noninvasive nature, and the high degree of spatial and temporal resolution attainable, light has become the tool of choice for remote manipulation of biological systems. For instance, photoactivatable green-fluorescent protein (GFP) has enabled numerous studies that track protein movement or mark specific cells in a population.^19^ Similarly, light controllable tetrazine ligations could open up applications such as cell surface engineering with single-cell precision and timed drug release.^20^ Recent efforts toward engineering light-sensitive dienophiles such as caged cyclopropenes^21^ and bicyclononynes^22^ have enabled ultraviolet light-induced tetrazine ligations with relatively modest reaction rates. However, ultraviolet light is toxic to live cells, particularly mammalian cells, which limits applications.^23^ *Trans-cyclooctenes,* the fastest reacting dienophiles, and caged “click to release” dienophiles have not yet been shown to be amenable to photocaging.^24^ In principle, light-triggered tetrazine formation would circumvent these issues. A visible light-triggered oxidation of air-stable 1,4-dihydrotetrazines to tetrazines using methylene blue as a photosensitizer has been developed.^25^ However, methylene blue is toxic and the use of a diffusible mediator limits spatiotemporal control.^26^ We hypothesized that direct activation of tetrazines using visible light would address these challenges and enable precision chemistry in biological systems.

To develop a light-triggered tetrazine ligation with high spatiotemporal precision, we explored whether a tetrazine precursor could be caged by a photocleavable protecting group (Fig. 1a). Dihydrotetrazine, a precursor for tetrazine, is unreactive to dienophiles^25^ and, if sufficiently electron-rich, spontaneously oxidized by air through reaction with oxygen.^24^ Therefore, we asked whether the secondary amines of dihydrotetrazine could be modified with visible light-cleavable protecting groups, such as nitrophenyl derivatives.^27^ Photocaging would prevent the oxidation of dihydrotetrazine to tetrazine, creating a compound that is inactive to cycloaddition with strained dienophiles. The caging group would be removed upon exposure to visible blue light, releasing dihydrotetrazine, which is spontaneously oxidized to the reactive tetrazine by air. Subsequently, the tetrazine would be able to react rapidly with a dienophile through inverse Diels-Alder cycloaddition. By directly activating the tetrazine, spatial control could be achieved, and the responsiveness to visible light would enable live cell applications.

**Fig. 1.**
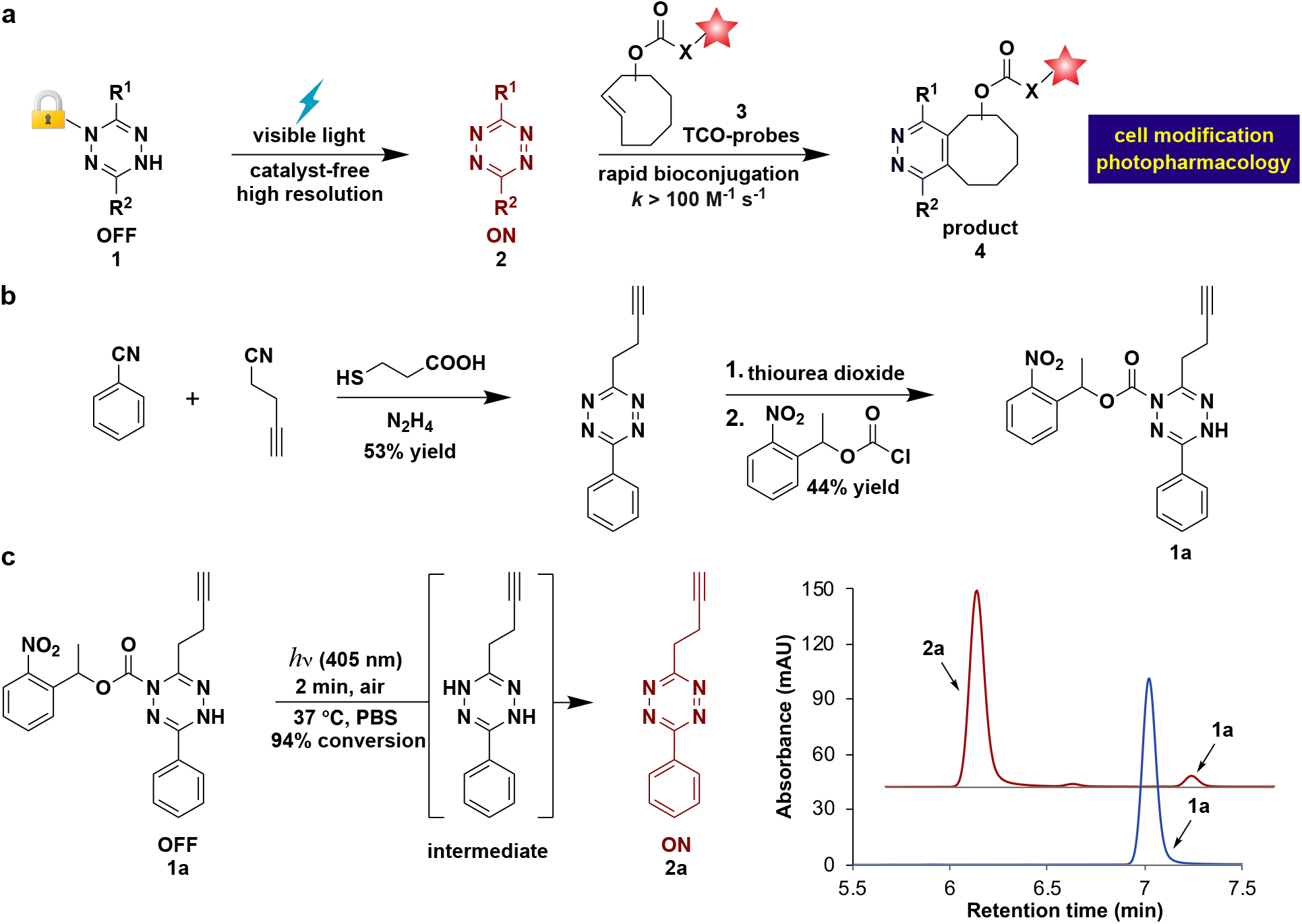
Light-controlled bioorthogonal tetrazine ligation in living cells. **a**, Visible light uncaging of a photoprotected dihydrotetrazine leads to formation of tetrazine which reacts rapidly with dienophiles such as *trans*-cyclooctene (TCO). Dienophiles could be appended to fluorescent probes (red star) or uncage prodrugs upon cycloaddition (“click to release”) enabling applications such as spatiotemporal modification of live cells or photopharmacology. **b,** Synthetic route to photocaged dihydrotetrazine **1a**. **c**, Light-activated formation of tetrazine **2a** from photocaged dihydrotetrazine **1a**. The reaction was carried out by irradiation of photocaged dihydrotetrazine **1a** (16 μM) with an LED light (405 nm, 18 W) in PBS solution (containing 0.1% DMSO) under open air at 37°C for 2 minutes. Samples were taken from the reaction mixture at different time points and examined by HPLC. Spectra before (blue) and after (red) LED irradiation (405 nm) for 2 minutes (absorbance monitored at 280 nm).

A photocaged dihydrotetrazine should be stable in aqueous solution and the uncaged dihydrotetrazine should rapidly be oxidized to tetrazine by air. We synthesized photocaged dihydrotetrazine **1a** from 3-(but-3-yn-1-yl)-6-phenyl-1,2,4,5-tetrazine, which could be converted to the corresponding dihydrotetrazine using the reductant thiourea dioxide (Fig. 1b and Fig. S1). 1-(2-Nitrophenyl)ethyl carbamate was chosen as the photocleavable functional group, due to its sensitivity to visible blue light,^28^ and biocompatibility.^29^ After reacting the dihydrotetrazine intermediate with the selected photocleavable group, the desired product 1-(2-nitrophenyl)ethyl 6-(but-3-yn-1-yl)-3-phenyl-1,2,4,5-tetrazine-1(4H)-carboxylate photocaged dihydrotetrazine **1a** was obtained (Fig. S1). Nuclear magnetic resonance (NMR) studies confirmed that the secondary amine adjacent to the alkyl group of dihydrotetrazine **1a** is functionalized (for details, see SI). We analysed light-triggered tetrazine formation from photocaged dihydrotetrazine **1a** (Fig. 1c). HPLC revealed that **1a** was nearly quantitatively decaged (94% conversion) when irradiated by visible blue light (405 nm, LED, 18 W) for 2 minutes in aqueous buffer (PBS containing 0.1% DMSO) at 37 *°C* (Fig. S2). During this time, the resulting dihydrotetrazine intermediate was spontaneously oxidized by air, generating the desired tetrazine **2a**. After demonstrating visible blue light-activated formation of tetrazine **2a,** we verified the stability of photocaged dihydrotetrazine **1a** in aqueous solution in the absence of irradiation. In phosphate-buffered saline (PBS) at 37 °C, degradation of **1a** was not observed over 1 day, and less than 1% of decomposition occurred after 4 days of incubation (Fig. S3). A major advantage of tetrazine ligations is the high rate of reaction with ring-strained *trans-cyclooctene* dienophiles. We measured a second-order rate constant of 101 ± 3 M^-1^ s^-1^ between **2a** and a model strained dienophile, *trans*-4-cycloocten-1-ol (TCO-OH), by monitoring the disappearance of the characteristic tetrazine visible absorption at 521 nm under pseudo first-order conditions (Fig. S4).^7^ The rapid rate constant encouraged us to move forward with exploring bioconjugation applications.

A drawback of tetrazines is their susceptibility to hydrolysis, particularly in the presence of nucleophiles and base.^30,31^ This feature has hampered applications of tetrazine ligations, particularly if tetrazines are required to be stable in physiological media for lengthy periods of time before cycloaddition, such as for pretargeted imaging, drug delivery applications, or in cases where early-stage modification of tetrazine is desired. For instance, although tetrazine ligations have been widely used to modify peptides, the tetrazine is typically introduced late-stage after peptide synthesis due to incompatibility with solid-phase peptide synthesis (SPPS) reaction conditions.^32^ Electron poor tetrazine-containing amino acids are not used in Fmoc solid-phase peptide synthesis (SPPS) due to their degradation during the standard repeated Fmoc deprotection conditions (e.g. 4-methylpiperidine/DMF).

Considering the stability of photocaged dihydrotetrazines, we asked if such groups could act as photoprotection moieties, masking latent tetrazines through harsh conditions that would normally be degradative. For example, in 4-methylpiperidine/DMF solution, 40% of tetrazine **2a** degrades in 30 minutes (Fig. S5). In contrast, photocaged dihydrotetrazine **1a** is stable in 4-methylpiperidine/DMF solution with no detectable degradation observed under the same reaction conditions. Therefore, we speculated that photocaged dihydrotetrazines would be tolerated during SPPS. To test this, we synthesized an unnatural Fmoc protected amino acid containing photocaged dihydrotetrazine, **1b** (Fig. S6). We employed **1b** in SPPS to obtain a five-amino-acid peptide **1c** in 50% yield (Fig. S7). After LED irradiation of photocaged dihydrotetrazine-peptide **1c** (10 μM) for 2 minutes in PBS at 37 °C, tetrazine-peptide **2c** was formed in 96% yield (Fig. 2a). Being able to directly use photoprotected tetrazine amino acids facilitates SPPS of peptides that contain multiple bioorthogonal handles, such as an azido group and tetrazine (Fig. S8), making it easier to site-specifically label peptides with multiple probes.

**Fig. 2.**
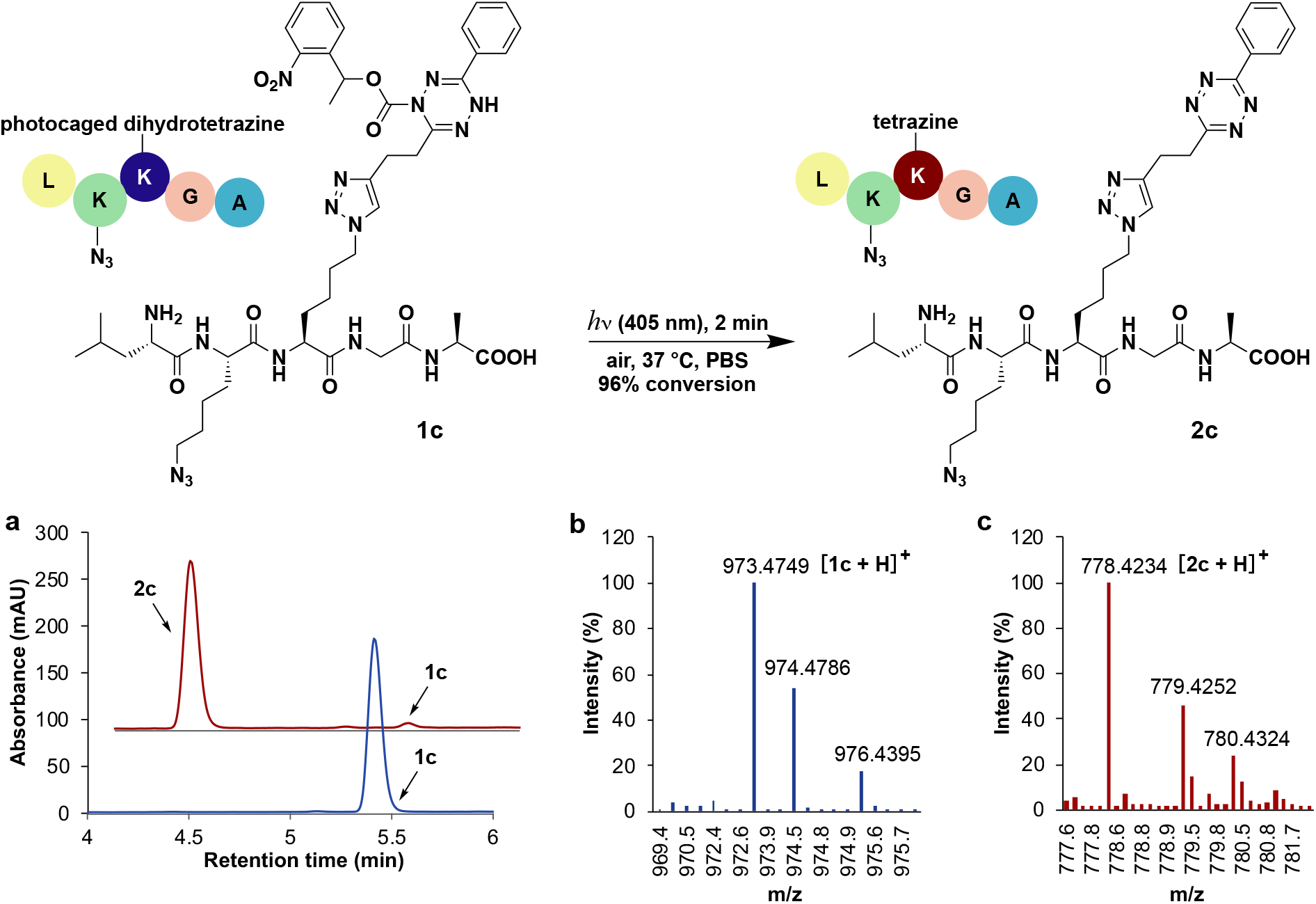
Early-stage functionalization of a peptide with a photocaged dihydrotetrazine group. Light-triggered formation of tetrazine-peptide **2c** from peptide **1c**, where a photocaged dihydrotetrazine amino acid was introduced during Fmoc solid phase peptide synthesis (SPPS). The reaction was carried out by irradiation of **1c** (10 μM) with LED light (405 nm, 18 W) in PBS solution (containing 0.2% DMSO and 0.2% DMF) under open air at 37 C for 2 minutes. **a,** HPLC/ELSD spectra of the peptide taken before (blue) and after (red) irradiation. **b**, HRMS spectrometry of photocaged-dihydrotetrazine-peptide **1c**. Expected mass 973.4751 Da, found mass 973.4749 Da. **c**, HRMS spectrometry of tetrazine-peptide **2c**. Expected mass 778.4220 Da, found mass 778.4234 Da.

Since our strategy utilizes visible light to directly generate tetrazines from their precursors, we asked whether photocaged dihydrotetrazines could spatiotemporally modify living cells, for instance by covalently labeling membrane lipids (Figure 3a). To remodel cell membranes with a photocaged dihydrotetrazine, we appended **1a** to a derivative of 1,2-dipalmitoyl-*sn*-glycero-3-phosphoethanolamine (DPPE) to form photocaged dihydrotetrazine-diacylphospholipid **1d** (Fig. 3b and Fig. S9). Upon irradiation with visible light (405 nm, LED, 18 W) **1d** reacted rapidly with a water-soluble *trans*-cyclooctene modified Alexa Fluor 488 dye (TCO-AF488) **3a** forming cycloaddition product **4a**, as determined by LC-MS (Fig. S10). To incorporate photocaged dihydrotetrazine onto cell membranes, adherent HeLa S3 cells were incubated with 60 nM photocaged dihydrotetrazine-diacylphospholipid **1d** in PBS solution (containing 0.1% DMSO) at 37 °C for 5 minutes. Excess **1d** was removed by washing cells with PBS solution. The washed cells were then incubated with 3 nM TCO-AF488 **3a** in PBS solution (containing 0.1% DMSO). To trigger in situ formation of tetrazine and the subsequent bioorthogonal tetrazine ligation, a single cell among a population of cells was selectively irradiated with a 405 nm laser (20 mW) for 20 seconds using a ZEISS 880 laser scanning microscope. Five minutes after the laser uncaging event, unreacted TCO-AF488 was removed by exchanging the solution with fresh cell culture medium. Fluorescence live-cell imaging was performed to reveal if tetrazine ligation had taken place on the membrane of the laser irradiated cell. Fluorescence labeling by AF488 was only observed on the membrane of the laser irradiated cell and not on adjacent cells, illustrating that precise spatiotemporal photo-activation of tetrazine ligation can be achieved using photocaged dihydrotetrazine-diacylphospholipid **1d** (Fig. 3c).

**Fig. 3.**
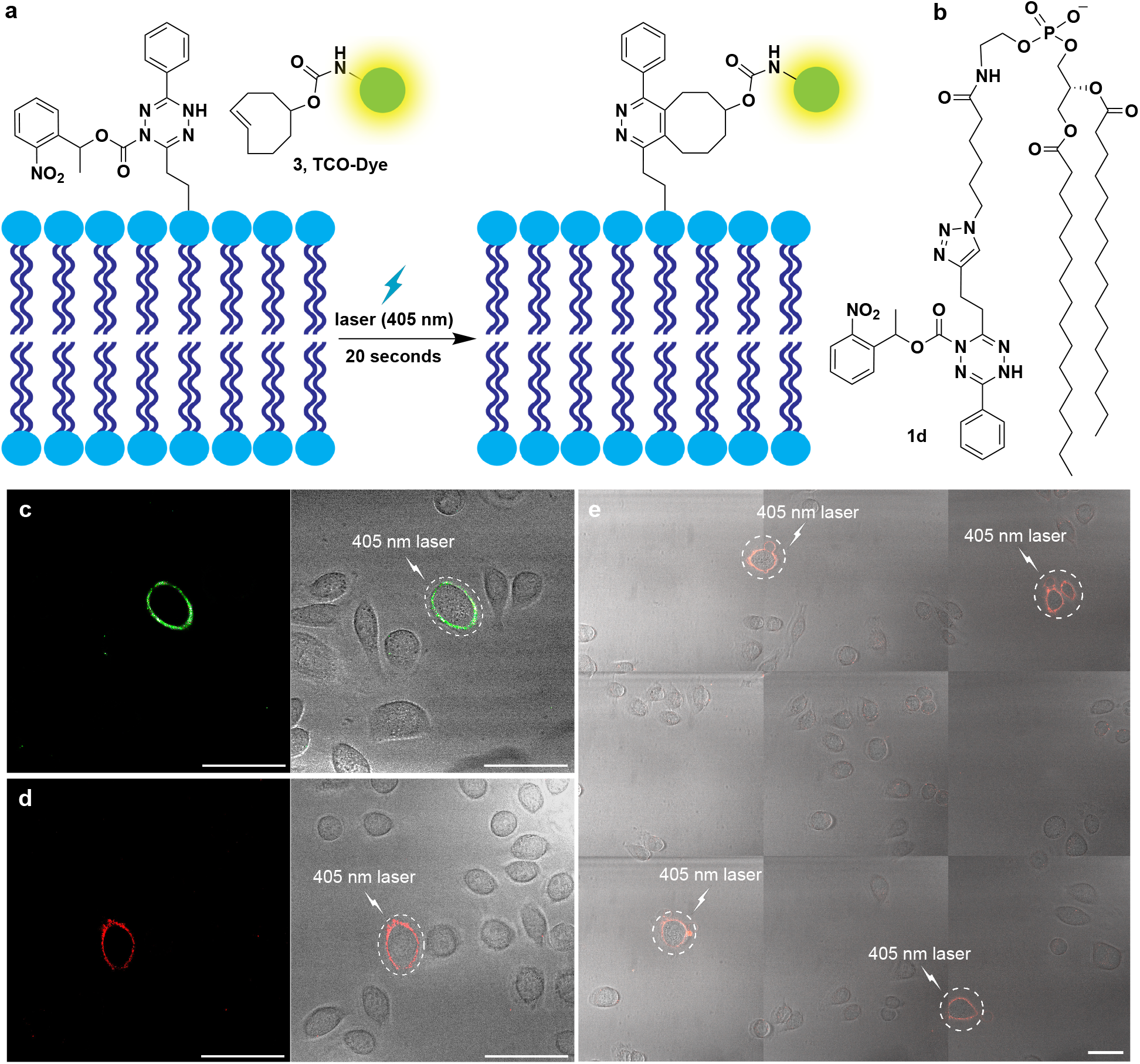
Single-cell remodeling of HeLa S3 cell membranes by photo-activation of tetrazine ligation. **a**, Cartoon depicting live cell photo-activation of tetrazine ligation on cellular membranes using photocaged dihydrotetrazine-diacylphospholipid **1d** and a *trans*-cyclooctene modified dye (TCO-Dye). **b**, Photocaged dihydrotetrazine-diacylphospholipid **1d**. **c**, Fluorescence live-cell labeling demonstrating single-cell photo-activation of tetrazine ligation on the cell membrane of a selected HeLa S3 cell. TCO-AF488 was used for tetrazine ligation. Fluorescence channel (AF488) shown on the left, and merged fluorescence and brightfield channels on the right. The area irradiated by the 405 nm laser is denoted by the white dashed circle. **d**, Spatiotemporal photo-activation of tetrazine ligation using TCO-AF568. Fluorescence channel (AF568) shown on the left, and merged fluorescence and brightfield channels on the right. The area irradiated by the 405 nm laser is denoted by the white dashed circle in the merged channel. **e**, Activation of four groups of cells at different locations inside a 0.75 mm by 0.75 mm square area. TCO-AF568 was used for the tetrazine ligation. Images taken from the merged fluorescence (AF568) and brightfield channels. The areas irradiated by the 405 nm laser are denoted by the white dashed circles. Scale bar: 50 μm.

To demonstrate the versatility of the light-activated single-cell manipulation, we also performed labeling with an alternative *trans*-cyclooctene modified dye, Alexa Fluor 568 dye (TCO-AF568) **3b** (Fig. 3d and Fig. S11). Additionally, our technique can be applied to other types of mammalian cells, for example, Hep 3B human liver cancer cells (Fig. S12). Finally, to test the robustness of spatial photo-activation, populations of 1-2 cells located at four different locations inside a 0.75 mm by 0.75 mm area were selectively laser irradiated at 405 nm to trigger tetrazine ligation, modifying the associated cell membranes (Fig. 3e). Fluorescence labeling was only observed on the laser irradiated cells, demonstrating that reliable spatiotemporal labeling of living cells can be achieved by light-activation of surface tetrazines.

An application of tetrazine ligation is the so called “click to release” strategy, which typically involves utilizing a dienophile to cage a bioactive molecule such as a drug.^24^ Upon reaction with tetrazine, tautomerization of the cycloadduct occurs leading to elimination and drug release.^6^ We speculated that combining light activated tetrazine formation with “click to release” strategies would facilitate the controlled release of therapeutics in the presence of living systems for photopharmacology. To couple light-activated tetrazine ligation with “click to release”, we synthesized a dienophile modified prodrug, *trans-cyclooctene* carbamate-caged doxorubicin **3c** (TCO-Dox) (Fig. S13), which liberates doxorubicin **5a** (Dox) after undergoing cycloaddition reaction with tetrazine (Fig. S14). Doxorubicin is an anticancer drug, and we therefore sought to use light to stimulate doxorubicin delivery to cancer cells through the dual activation process (Fig. 4a), triggering apoptosis. We first tested the dual activation process by irradiating a reaction mixture consisting of photocaged dihydrotetrazine **1a** (8 μM) and TCO-Dox **3c** (5.5 μM) with LED light (405 nm, 18 W) for 2 minutes in PBS solution (containing 0.1% DMSO) at 37 °C. This was followed by incubation at 37 °C for 24 hours. Samples from the reaction mixture were taken at different time points, and analysis by HPLC-MS showed that 91% of TCO-Dox **3c** was converted to doxorubicin **5a** after 24 hours (Fig. S15). In the absence of irradiation, the mixture of photocaged dihydrotetrazine **1a** (8 μM) and TCO-Dox **3c** (5.5 μM) showed no reaction (Fig. S16).

**Fig. 4.**
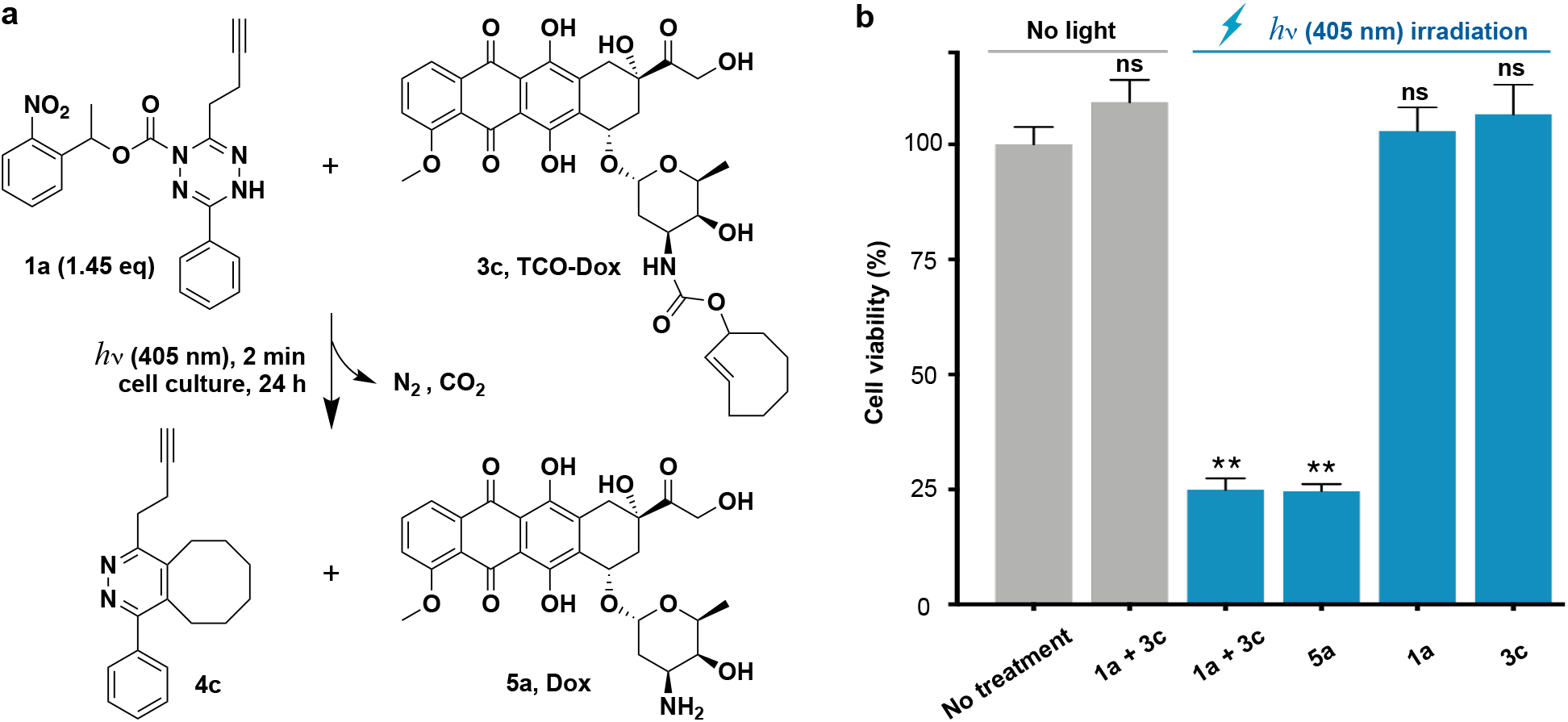
Light-activated tetrazine prodrug therapy in Hep 3B cancer cells. **a**, Application of light-controlled tetrazine ligation to release the chemotherapeutic doxorubicin in the presence of living Hep 3B cancer cells. Dox, doxorubicin. **b**, Cell viability of Hep 3B cancer cells after treatments with photocaged dihydrotetrazine **1a** (8 μM), TCO-Dox **3c** (5.5 μM), and Dox **5a** (5.5 μM) with or without irradiation by LED light (405 nm, 18 W) for 2 minutes, followed by incubation at 37 °C for 24 hours. Error bars indicate standard error of mean (SEM), which are measured from 3 replicates. Statistically significant differences in cell viability between no treatment and other means are indicated: *\*\*P* < 0.01. ns, not significant.

Based on the positive results obtained, we tested whether light-triggered drug delivery could be carried out in the presence of live cancer cells (Fig. 4b). When Hep 3B human liver cancer cells were treated with a mixture of photocaged dihydrotetrazine **1a** (8 μM) and TCO-Dox **3c** (5.5 μM), no decrease in cell viability was observed compared to untreated cells. However, upon irradiation with LED light (405 nm, 18W) for 2 minutes, followed by incubation for 24 hours, a 75.1 ± 3 % reduction of cell viability was observed. The loss of cell viability was similar to that observed when cells were directly treated with Dox **5a** (5.5 μM) under the same conditions. There was no influence on cell viability when cells were irradiated with LED light for 2 minutes in the presence of either photocaged dihydrotetrazine **1a** (8 μM) or TCO-Dox **3c** (5.5 μM) alone. These results demonstrate that photocaged dihydrotetrazines can be utilized for the light-triggered release of bioactive compounds, such as chemotherapeutics, in the presence of living cells.

Here we have demonstrated a methodology for the photoactivation of tetrazines that enables biomolecular labeling, spatiotemporal modification of live-cell membranes with single-cell precision, and photopharmacology when combined with “click to release” strategies. Tetrazine instability is a well-recognized obstacle to their use, and we have found that photocaged tetrazine precursors are highly stable, even in the presence of strong bases which rapidly degrade tetrazines. Given the stability of photocaged dihydrotetrazines, we expect they will find broad application as a general tetrazine protecting group. Photocaged dihydrotetrazines would be especially useful in conditions known to degrade tetrazines, such as those encountered during the installation of ^18^F radionuclides for PET imaging^30^ or for live-cell pulse-chase experiments where tetrazine reactivity would be required to be maintained for an arbitrary amount of time before reaction.^33^ Indeed, preliminary results indicate that photocaged dihydrotetrazines, unlike tetrazines, are very stable under the conditions typically used for fluorination (Fig S17). Since our method directly activates tetrazine precursors, high spatiotemporal precision is achievable. By modifying phospholipids on cell surfaces, we showed that single-cell activation is feasible. The technique could enable monitoring of lipid trafficking and dynamics in live cells by controlling where and when caged tetrazines on lipids are activated and following their transport by post-labeling with dienophile modified fluorophores.^34^ Light activated release of the chemotherapeutic doxorubicin was carried out by combining photoactivation of tetrazine formation with “click to release” strategies. The photocaged tetrazine precursor and light alone showed negligible toxicity, demonstrating the biocompatibility of the technique. Such optically controlled drug release may have practical application in image guided surgery and photodynamic therapy.^35^ Future studies will explore alternative caging groups that activate in response to longer wavelengths of light, enabling multiplexing and facilitating in vivo studies.^27^ The basic concept we present might also be extended to other amine caging functionalities capable of masking dihydrotetrazines, allowing rapid biorthogonal ligation in response to additional stimuli such as enzymatic activity, pH, or the presence of metal complexes.^36,37^

## Supporting information

Experimental materials and methods

## Acknowledgments

Financial support for this work is provided by the National Institutes of Health (DP2DK111801, R01GM123285 and T32CA009523-33). We thank Prof. Itay Budin and Dr. Guy Riddihough for reviewing this manuscript and providing helpful comments.

## Author contributions

L. L. and N. K. D. conceived the project. L. L. designed and performed the synthetic experiments. L. L., and D. Z. performed microscopy experiments. L. L., and M. J. performed cell experiments. L. L., D. Z., M. J., and N. K. D. analyzed the data. L. L., and N. K. D. wrote the manuscript.

## References

1. Saxon, E. & Bertozzi, C. R. Cell surface engineering by a modified staudinger reaction. Science. 287, 2007–2010 (2000).

2. Prescher, J. A., Dube, D. H. & Bertozzi, C. R. Chemical remodelling of cell surfaces in living animals. Nature. 430, 873–877 (2004).

3. Devaraj, N. K. The future of bioorthogonal chemistry. ACS Cent. Sci. 4, 952–959 (2018).

4. Oliveira, B. L., Guo, Z. & Bernardes, G. J. L. Inverse electron demand Diels-Alder reactions in chemical biology. Chem. Soc. Rev. 46, 4895–4950 (2017).

5. Nguyen, S. S. & Prescher, J. A. Developing bioorthogonal probes to span a spectrum of reactivities. Nat. Rev. Chem. 4, 476–489 (2020).

6. Carlson, J. C. T., Mikula, H. & Weissleder, R. Unraveling tetrazine-triggered bioorthogonal elimination enables chemical tools for ultrafast release and universal cleavage. J. Am. Chem. Soc. 140, 3603–3612 (2018).

7. Versteegen, R. M., ten Hoeve, W., Rossin, R., de Geus, M. A. R., Janssen, H. M. & Robillard, M. S. Click-to-release from *trans*-cyclooctenes: mechanistic insights and expansion of scope from established carbamate to remarkable ether cleavage. Angew. Chem. Int. Ed. 57, 10494–10499 (2018).

8. Czuban, M., Srinivasan, S., Yee, N. A., Agustin, E., Koliszak, A., Miller, E., Khan, I., Quinones, I., Noory, H., Motola, C., Volkmer, R., Di Luca, M., Trampuz, A., Royzen, M. & Mejia Oneto, J. M. Bio-orthogonal chemistry and reloadable biomaterial enable local activation of antibiotic prodrugs and enhance treatments against staphylococcus aureus infections. ACS Cent. Sci. 4, 1624–1632 (2018).

9. Wu, K., Yee, N., Srinivasan, S., Mahmoodi, A., Zakharian, M., Mejia Oneto, J. M. & Royzen, M. Click activated protodrugs against cancer increase the therapeutic potential of chemotherapy through local capture and activation. Preprint at http://doi.org/10.26434/chemrxiv.13087715.v1 (2020).

10. Blackman, M. L., Royzen, M. & Fox, J. M. Rapid tetrazine ligation: fast bioconjugation based on inverse-electron-demand Diels-Alder reactivity. J. Am. Chem. Soc. 130, 13518–13519 (2008).

11. Devaraj, N. K., Weissleder, R. & Hilderbrand, S. A. Tetrazine-based cycloadditions: application to pretargeted live cell imaging. Bioconjugate Chem. 19, 2297–2299 (2008).

12. Elliott, T. S., Townsley, F. M., Bianco, A., Ernst, R. J., Sachdeva, A., Elsässer, S. J., Davis, L., Lang, K., Pisa, R., Greiss, S., Lilley, K. S. & Chin, J. W. Proteome labeling and protein identification in specific tissues and at specific developmental stages in an animal. Nat. Biotechnol. 32, 465–472 (2014).

13. Dong, J., Jan, Y. J., Cheng, J., Zhang, R. Y., Meng, M., Smalley, M., Chen, P. J., Tang, X., Tseng, P., Bao, L., Huang, T. Y., Zhou, D., Liu, Y., Chai, X., Zhang, H., Zhou, A., Agopian, V. G., Posadas, E. M., Shyue, J. J., Jonas, S. J., Weiss, P. S., Li, M., Zheng, G., Yu, H. H., Zhao, M., Tseng, H. R., & Zhu, Y. Covalent chemistry on nanostructured substrates enables noninvasive quantification of gene rearrangements in circulating tumor cells. Sci. Adv. 5, eaav9186 (2019).

14. Liang, D., Wu, K., Tei, R., Bumpus, T. W., Ye, J. & Baskin, J. M. A real-time, click chemistry imaging approach reveals stimulus-specific subcellular locations of phospholipase D activity. Proc. Natl. Acad. Sci. USA 116, 15453–15462 (2019).

15. Agarwal, P., Beahm, B. J., Shieh, P. & Bertozzi, C. R. Systemic fluorescence imaging of zebrafish glycans with bioorthogonal chemistry. Angew. Chem. Int. Ed. 54, 11504–11510 (2015).

16. Li, H., Conde, J., Guerreiro, A. & Bernardes, G. J. L. Tetrazine carbon nanotubesfor pretargeted in vivo “click-to-release” bioorthogonal tumour imaging. Angew. Chem. Int. Ed. 59, 16023–16032 (2020).

17. Ji, X., Pan, Z., Yu, B., De La Cruz, L. K., Zheng, Y., Ke, B. & Wang, B. Click and release: bioorthogonal approaches to “on-demand” activation of prodrugs. Chem. Soc. Rev. 48, 1077–1094 (2019).

18. Li, J., Jia, S., & Chen, P. R. Diels-Alder reaction-triggered bioorthogonal protein decaging in living cells. Nat. Chem. Biol. 10, 1003–1005 (2014).

19. Patterson, G. H. & Lippincott-Schwartz, J. A photoactivatable GFP for selective photolabeling of proteins and cells. Science. 297, 1873–1887 (2002).

20. Kumar, G. S. & Lin, Q. Light-triggered click chemistry. Chem. Rev. http://doi.org/10.1021/acs.chemrev.0c00799 (2020).

21. Kumar, P., Jiang, T., Li, S., Zainul, O. & Laughlin, S. T. Caged cyclopropenes for controlling bioorthogonal reactivity. Org. Biomol. Chem. 16, 4081–4085 (2018).

22. Mayer, S. V., Murnauer, A., von Wrisberg, M. K., Jokisch, M. L. & Lang, K. Photo-induced and rapid labeling of tetrazine-bearing proteins via cyclopropenone-caged bicyclononynes. Angew. Chem. Int. Ed. 58, 15876–15882 (2019).

23. Hockberger, P. E. A history of ultraviolet photobiology for humans, animals and microorganisms. Photochem. Photobiol. 76, 561–579 (2002).

24. Versteegen, R. M., Rossin, R., ten Hoeve, W., Janssen, H. M. & Robillard, M. S. Click to release: instantaneous doxorubicin elimination upon tetrazine ligation. Angew. Chem. Int. Ed. 52, 14112–14116 (2013).

25. Zhang, H., Trout, W. S., Liu, S., Andrade, G. A., Hudson, D. A., Scinto, S. L., Dicker, K. T., Li, Y., Lazouski, N., Rosenthal, J., Thorpe, C., Jia, X. & Fox, J. M. Rapid bioorthogonal chemistry turn-on through enzymatic or long wavelength photocatalytic activation of tetrazine ligation. J. Am. Chem. Soc. 138, 5978–5983 (2016).

26. da Costa S. R., da Costa Monteiro M., da Silva Júnior F. M. R. & Sandrini J. Z. Methylene blue toxicity in zebrafish cell line is dependent on light exposure. Cell Biol. Int. 40, 895–905 (2016).

27. Hansen, M. J., Velema, W. A., Lerch, M. M., Szymanski, W. & Feringa, B. L. Wavelength-selective cleavage of photoprotecting groups: strategies and applications in dynamic systems. Chem. Soc. Rev. 44, 3358–3377 (2015).

28. Corrie, J. E. T., Barth, A., Munasinghe, V. R. N., Trentham, D. R. & Hutter, M. C. Photolytic cleavage of 1-(2-nitrophenyl)ethyl ethers involves two parallel pathways and product release is rate-limited by decomposition of a common hemiacetal intermediate. J. Am. Chem. Soc. 125, 8546–8554 (2003).

29. Zhao, Y. R., Zheng, Q., Dakin, K., Xu, K., Martinez, M. L. & Li, W. H. New caged coumarin fluorophores with extraordinary uncaging cross sections suitable for biological imaging applications. J. Am. Chem. Soc. 126, 4653–4663 (2004).

30. Li, Z., Cai, H., Hassink, M., Blackman, M. L., Brown, R. C. D., Conti, P. S. & Fox, J. M. Tetrazine-*trans*-cyclooctene ligation for the rapid construction of ^18^F labeled probes. Chem. Commun. 46, 8043–8045 (2010).

31. Selvaraj, R., Liu, S., Hassink, M., Huang, C. W., Yap, L. P., Park, R., Fox, J. M., Li, Z. & Conti, P. S. Tetrazine-*trans*-cyclooctene ligation for the rapid construction of integrin αvβ3 targeted PET tracer based on a cyclic RGD peptide. Bioorg. Med. Chem. Lett. 21, 5011–5014 (2011).

32. Pagel, M. Inverse electron demand Diels-Alder (IEDDA) reactions in peptide chemistry. J. Pept. Sci. 25, e3141. (2019).

33. Willis, J. C. W. & Chin, J. W. Mutually orthogonal pyrrolysyl-tRNA synthetase/tRNA pairs. Nat. Chem. 10, 831–837 (2018).

34. Tamura, T., Fujisawa, A., Tsuchiya, M., Shen, Y., Nagao, K., Kawano, S., Tamura, Y., Endo, T., Umeda, M. & Hamachi, I. Organelle membrane-specific chemical labeling and dynamic imaging in living cells. Nat. Chem. Biol. 16, 1361–1367 (2020).

35. Kwiatkowski, S., Knap, B., Przystupski, D., Saczko, J., Kędzierska, E., Knap-Czop, K., Kotlińska, J., Michel, O., Kotowski, K. & Kulbacka, J. Photodynamic therapy-mechanisms, photosensitizers and combinations. Biomed. Pharmacother. 106, 1098–1107 (2013).

36. Cao, Z., Li, W., Liu, R., Li, X., Li, H., Liu, L., Chen, Y., Lv, C. & Liu, Y. pH- and enzyme-triggered drug release as an important process in the design of anti-tumor drug delivery systems. Biomed. Pharmacother. 118, 109340 (2019).

37. Li, J., Yu, J., Zhao, J., Wang, J., Zheng, S., Lin, S., Chen, L., Yang, M., Jia, S., Zhang, X. & Chen, P. R. Palladium-triggered deprotection chemistry for protein activation in living cells. Nat. Chem. 6, 352–361 (2014).

