## Supplementary material for "Light-activated tetrazines enable live-cell spatiotemporal control of bioorthogonal reactions": Experimental materials and methods

#### This PDF file includes:

Materials and Methods
Supplementary Text
Supplementary Figures 1–17
Copies of Mass and NMR Spectra
References

#### Table of Contents


| | | |
|---|---|---|
| | General Information | S3 |
| 1. | Synthesis of Photocaged Dihydrotetrazine **1a** | S4 |
| 2. | Light-Activated Formation of Tetrazine **2a** | S5 |
| 3. | Kinetic Study of the Reaction between Tetrazine **2a** and *Trans*-4-cycloocten-1-ol | S7 |
| 4. | Application of Photocaged Dihydrotetrazine **1** in Solid-Phase Peptide Synthesis | S8 |
| 5. | Spatiotemporal Live-Cell Labeling with Alexa Fluor Dyes | S13 |
| 6. | Delivery of Doxorubicin by Light-Activated Tetrazine Ligation | S17 |
| 7. | The Stabilities of **1a** and **2a** under Fluorination Reaction Conditions | S26 |
| 8. | NMR Spectra | S27 |
| 9. | References | S34 |

#### General information

Chemicals were purchased as reagent grade and used without further purification except as indicated below. Solvents ($CH_2Cl_2$, $CHCl_3$, MeOH, DMF, THF, $H_2O$) were purchased from commercial suppliers. Axial TCO-PNP was purchased from SiChem GmbH. Light-irradiation was carried out with an LED light (405 nm, 18 W), which was purchased from HepatoChem Inc (P206-18-1). Thin-layer chromatography (TLC) was performed using silica gel pre-coated plastic sheets (Polygram SIL G/$UV_{254}$, 0.2 mm, with fluorescent indicator; Macherey-Nagel), which were visualized with a UV lamp (254 nm), or Potassium Permanganate stain (1.5g of $KMnO_4$, 10g $K_2CO_3$, and 1.25mL 10% NaOH in 200 mL $H_2O$). Column chromatography was carried out using Merck silica gel (60 Å, 230−400 mesh, particle size 0.040−0.063 mm) using technical grade solvents. Nuclear magnetic resonance (NMR) spectra were recorded on a Bruker VX-500, or AVIII-600 NMR spectrometer in a suitable deuterated solvent. Chemical shifts are reported with tetramethylsilane (TMS) serving as the internal standard for all nuclides. The resonance multiplicity is described as s (singlet), d (doublet), t (triplet), q (quadruplet), m (multiplet), and br (broad). All spectra were recorded at 298 K unless otherwise noted, processed with MestReNova 12.0.0 suite, and coupling constants are reported as observed. An Agilent 6230 time-of-flight mass spectrometer (TOF-MS) with Jet Stream electrospray ionization source (ESI) was used for high-resolution mass spectrometry (HR-MS) analysis. The Jet Stream ESI source was operated under positive ion mode with the following parameters: VCap: 3500 V; fragmentor voltage: 160 V; nozzle voltage: 500 V; drying gas temperature: 325 °C, sheath gas temperature: 325 °C, drying gas flow rate: 7.0 L/min; sheath gas flow rate: 10 L/min; nebulizer pressure: 40 psi. Liquid chromatography-mass spectrometry (LC-MS) was performed on an Agilent 6120 quadrupole LC-MS equipped with a diode array detector (DAD), an evaporative light scattering detector (ELSD), and a mass spectrometer (Agilent 1260 infinity), using an Eclipse Plus C8 analytical column. Spinning-disk confocal microscopy images were acquired on a Yokogawa spinning-disk system (Yokogawa, Japan) built around an Axio Observer Z1 motorized inverted microscope (Carl Zeiss S-3 Microscopy GmbH, Germany) with a 20x, 0.8 NA objective or a 63x, 1.40 NA oil immersion objective. Images were captured with an ORCA-Flash4.0 V2 Digital CMOS camera (Hamamatsu, Japan) using ZEN Blue imaging software (Carl Zeiss Microscopy GmbH, Germany). The fluorophores were excited with diode lasers (405 nm-20 mW, 488 nm-30 mW, and 561 nm-20 mW). A condenser/objective with a phase stop of Ph3 was used to obtain the phase-contrast images. Absorbance measurements were performed on a Tecan infinite F200 plate reader instrument.

#### 1. Synthesis of Photocaged Dihydrotetrazine 1a

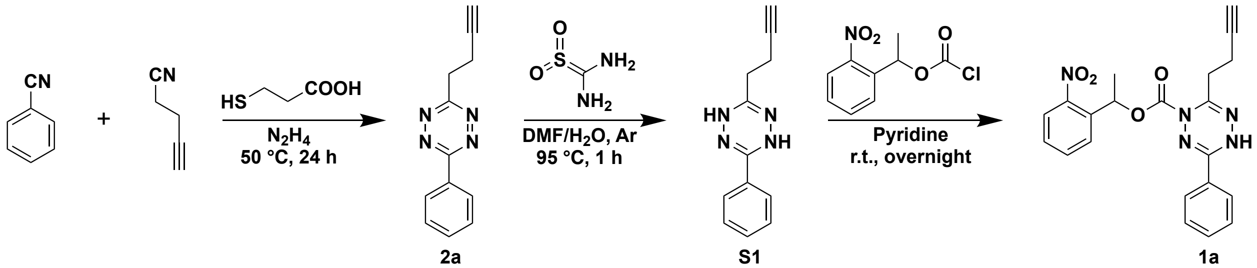


**Fig. 1. Synthesis of photocaged dihydrotetrazine 1a.**

A mixture of benzonitrile (202 μL, 2 mmol), 4-pentynenitrile (720 μL, 8 mmol), and 3-mercaptopropionic acid (88 μL, 1 mmol) was cooled to 0 °C under argon. Anhydrous hydrazine (1.55 mL, 32 mmol) was added dropwise to the mixture. The reaction mixture was stirred in an oil bath at 50 °C for 24 hours. Upon completion, the reaction solution was cooled with ice water. A solution of sodium nitrite (2 g, 30 mmol) in ice water was slowly added into the reaction mixture, followed by slow addition of 1M HCl. Addition of 1M HCl continued until gas evolution ceased. Then, the reaction mixture was extracted with $CH_2Cl_2$ and washed with a saturated solution of NaCl in water. The extract was combined, dried over $Na_2SO_4$ and concentrated under reduced pressure. The residue was purified by column chromatography on silica gel using 30%–50% $CH_2Cl_2$/Hexane as the eluents yielding the title compound **2a** as a pink solid (224 mg, 53%).[1] In a sealed flask, the reductant thiourea dioxide (32 mg, 0.3 mmol) was added to a solution of tetrazine **2a** (42 mg, 0.2 mmol) in 1.5 mL of dimethylformamide (DMF)/$H_2O$ (v/v = 10/1) at room temperature under argon. The reaction mixture was stirred in an oil bath at 95 °C for 1 hour.[2] Upon completion, the color of the reaction mixture changed from pink to light yellow. The reaction solvent was removed under reduced pressure and the residue was dried under high vacuum overnight, resulting in a powder containing dihydrotetrazine **S1**, which was transferred to a sealed flask under argon and directly used for next step. Under argon, 3.5 mL of pyridine was added, followed by slow addition of a solution of 1-(2-nitrophenyl)ethyl carbonochloridate[3] (115 mg, 0.5 mmol) in toluene (0.6 mL) at room temperature. The reaction mixture was stirred at room temperature overnight. Upon completion, the reaction mixture was concentrated under reduced pressure. The residue was purified by column chromatography on silica gel using 50%–100% $CH_2Cl_2$/Hexane to 15% MeOH/$CH_2Cl_2$ as the eluents yielding the title compound **1a** as a pale-yellow solid (36 mg, 44%).

**1-(2-Nitrophenyl)ethyl 6-(but-3-yn-1-yl)-3-phenyl-1,2,4,5-tetrazine-1(4H)-carboxylate (1a)**

**$^1$H NMR** (600 MHz, $(CD_3)_2SO$): δ 10.28 (s, 1 H), 8.01 (dd, $J$ =12.0 Hz, $J$ = 6.0 Hz, 1 H), 7.87–7.86 (m, 2 H), 7.82 (td, $J$ = 12.0 Hz, $J$ = 6.0 Hz, 1 H), 7.78 (dd, $J$ = 12.0 Hz, $J$ = 6.0 Hz, 1 H), 7.61–7.57 (m, 2 H), 7.53–7.51 (m, 2 H), 6.22 (q, $J$ = 6.0 Hz, 1 H), 2.82 (t, $J$ = 6.0 Hz, 2 H), 2.79 (t, $J$ = 6.0 Hz, 1 H), 2.37 (qd, $J$ = 12.0 Hz, $J$ = 6.0 Hz, 2 H), 1.69 (q, $J$ = 6.0 Hz, 3 H).

**$^{13}$C NMR** (151 MHz, $(CD_3)_2SO$): δ 155.66, 149.31, 147.45, 142.01, 136.74, 134.20, 131.67, 129.15, 128.78, 128.39, 127.34, 126.93, 124.30, 82.86, 71.93, 69.50, 29.71, 21.60, 15.26.

**HRMS** *m/z* (ESI): calcd. for $C_{21}H_{20}N_5O_4$ $[M+H]^+$: 406.1510; found: 406.1507.

**3-(But-3-yn-1-yl)-6-phenyl-1,2,4,5-tetrazine (2a)**

**$^1$H NMR** (500 MHz, $CD_3OD$): δ 8.62 (td, $J$ = 10.0 Hz, $J$ = 5.0 Hz, 2 H), 7.66–7.59 (m, 3 H), 3.61 (dt, $J$ = 10.0 Hz, $J$ = 5.0 Hz, 2 H), 2.95 (td, $J$ = 10.0 Hz, $J$ = 5.0 Hz, 2 H), 2.0 (t, $J$ = 5.0 Hz, 1 H).

$^{13}$**C NMR** (126 MHz, $CD_3OD$): δ 168.31, 164.59, 132.92, 131.75, 129.44, 128.18, 81.97, 70.30, 33.88, 17.16.
**HRMS** *m/z* (ESI): calcd. for $C_{12}H_{11}N_4$ $[M+H]^+$: 211.0978; found: 211.0977.

#### 2. Light-Activated Formation of Tetrazine 2a

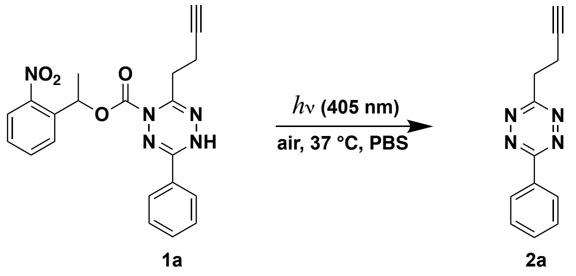


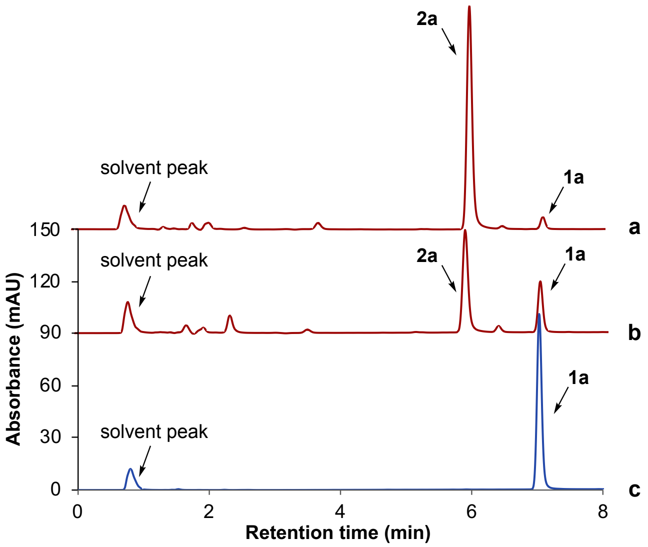


**Fig. 2. Light-triggered formation of tetrazine 2a from photocaged dihydrotetrazine 1a in phosphate-buffered saline (PBS).** The reaction was carried out by irradiation of photocaged dihydrotetrazine **1a** (16 μM) by LED light (405 nm, 18W) in PBS solution (containing 0.1% DMSO) open to air at 37 °C. Samples taken from the resulting reaction mixture at different time points were examined and analyzed by LC-MS. **a**, The reaction mixture was irradiated by LED light (405 nm, 18W) for 2 minutes. **b**, The reaction mixture was irradiated by LED light (405 nm, 18W) for 1 minute. **c**, No irradiation of the reaction mixture was carried out.

HPLC analysis was carried out on an Eclipse Plus C8 analytical column with Phase A/Phase B gradients [Phase A: MeOH with 0.1% trifluoroacetic acid, Phase B: $H_2O$ with 0.1% trifluoroacetic acid]. 50% Phase A in Phase B, 1 minute, and 50%–70% Phase A in Phase B, 4 minutes, then 70%–90% Phase A in Phase B, 3 minutes. The absorbance was monitored at 280 nm.
As shown in Fig. 2, 65% of photocaged dihydrotetrazine **1a** was decaged after 1 minute of irradiation, and 94% of photocaged dihydrotetrazine **1a** was decaged and converted to tetrazine **2a** after 2 minutes of irradiation.

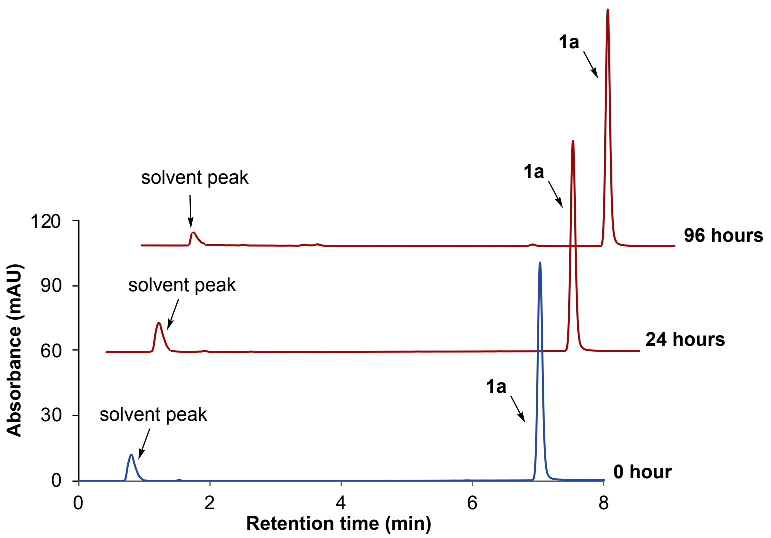


**Fig. 3. HPLC analysis of the stability of photocaged dihydrotetrazine 1a in PBS solution.** In the absence of light, 16 μM **1a** in PBS solution (containing 0.1% DMSO) was incubated at 37 °C in a sealed flask for 96 hours. Samples of the reaction mixture were taken at different time points and examined by LC-MS.

HPLC analysis was carried out on an Eclipse Plus C8 analytical column with Phase A/Phase B gradients [Phase A: MeOH with 0.1% trifluoroacetic acid, Phase B: $H_2O$ with 0.1% trifluoroacetic acid]. 50% Phase A in Phase B, 1 minute, and 50%–70% Phase A in Phase B, 4 minutes, then 70%–90% Phase A in Phase B, 3 minutes. The absorbance was monitored at 280 nm.

As shown in Fig. 3, there is no observed degradation of photocaged dihydrotetrazine **1a** in PBS solution at 37 °C for 24 hours. Less than 1% of dihydrotetrazine **1a** was degraded after 96 hours.

#### 3. Kinetic Study of the Reaction between Tetrazine 2a and *Trans*-4-cycloocten-1-ol (TCO-OH)

The second order reaction rate between tetrazine **2a** and *trans*-4-cycloocten-1-ol (TCO-OH) was estimated under pseudo first order conditions adapting a previously reported procedure.[4] Experiments were performed in PBS solution (containing 10% DMSO). 10 µL of a 1.3 mM solution of TCO-OH in PBS solution (containing 10% DMSO) was added to solutions of tetrazine **2a** to reach initial concentrations of 13 µM TCO-OH and 265 µM to 305 µM tetrazine **2a**. The decrease in the absorbance of tetrazine **2a** at 521 nm was monitored over time. Controls monitoring tetrazine absorbance in solvent alone strongly support that the loss of tetrazine during reaction is due solely to the reaction with TCO-OH, and thus the rate at which TCO-OH is consumed was obtained. The pseudo first order reaction rate constant $k_1$ was determined from the slope of a plot of -ln(C/C$_0$) versus time. The second order reaction rate constant $k_2$ was determined from the slope of a plot of the observed pseudo first order rate constant $k_1$ versus different concentrations of **2a**. All experiments were performed at 20 °C. Experiments were performed in 3 replicates.

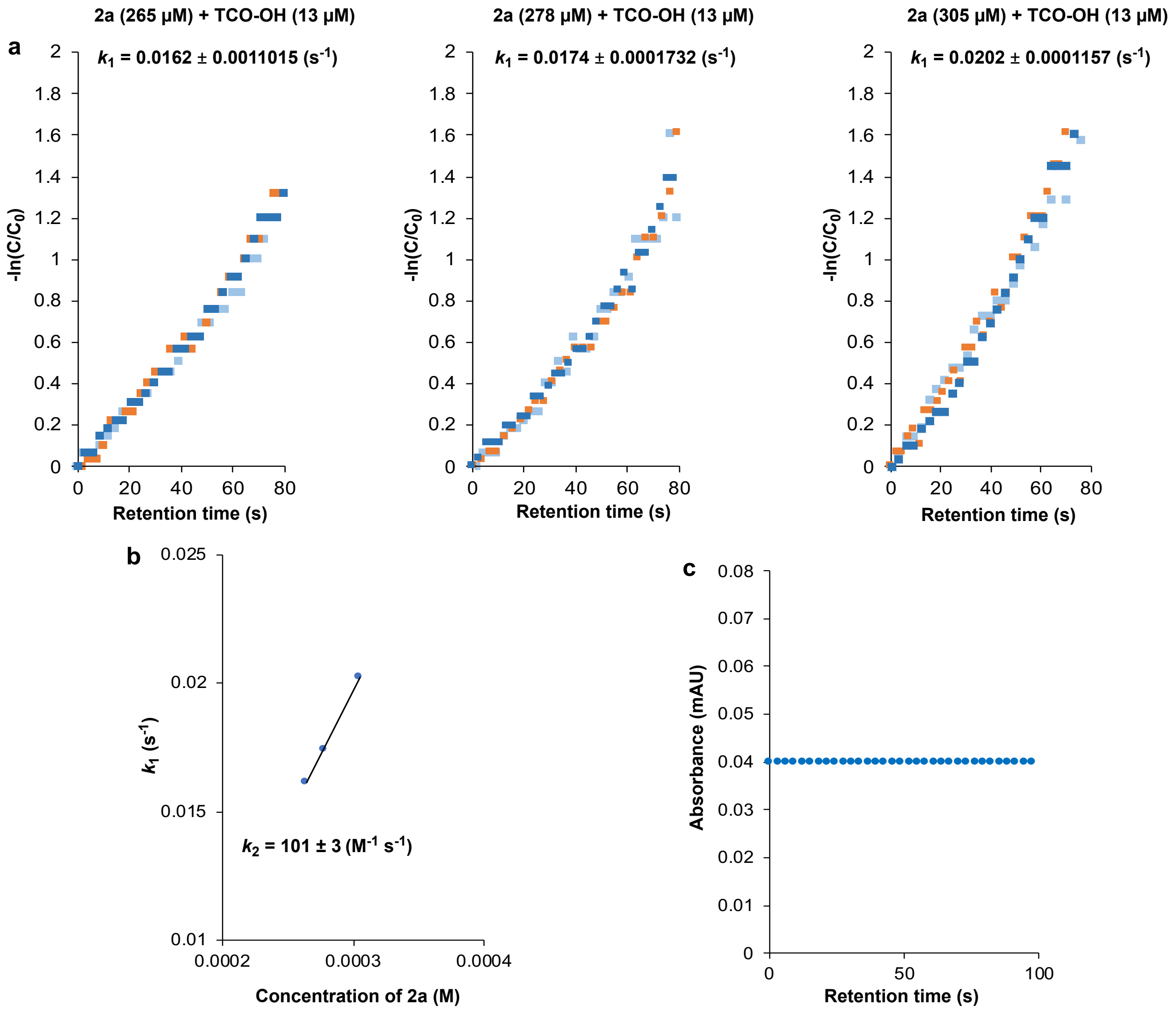

**Fig. 4. Kinetic study of the reaction between tetrazines 2a and *trans*-4-cyclooocten-1-ol (TCO-OH). a,** Determination of the pseudo first order reaction rate constant $k_1$ between TCO-OH (13 μM) and tetrazine **2a** (265 μM to 305 μM). **b,** Determination of second order reaction rate constant $k_2$. **c,** The absorbance of tetrazine **2a** (192 μM) at 521 nm in PBS solution (containing 10% DMSO) was followed over time.

#### 4. Application of Photocaged Dihydrotetrazine 1 in Solid-Phase Peptide Synthesis

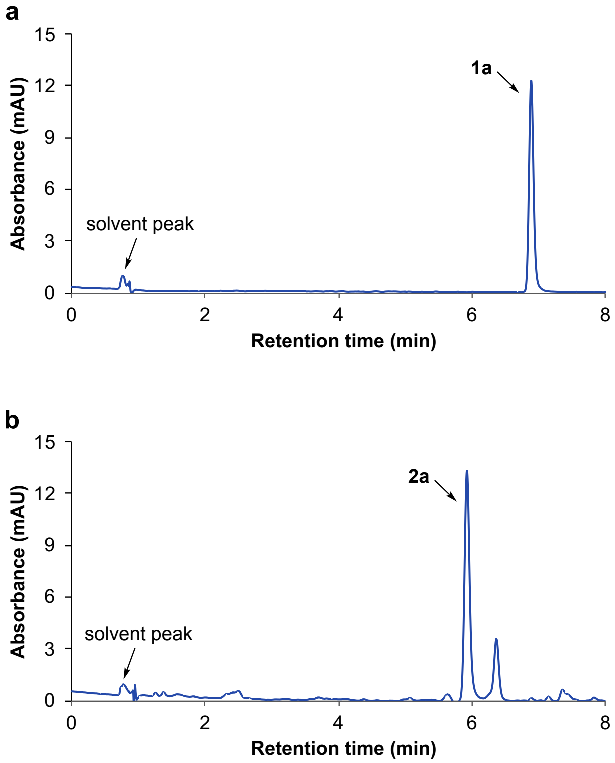


**Fig. 5. The stability of photocaged dihydrotetrazine 1a and tetrazine 2a in 4-methylpiperidine solution. a,** In the absence of light, 5 mM **1a** in a solution of 20% (w/w) 4-methylpiperidine/DMF, was incubated at 37 °C for 30 minutes. **b,** In the absence of light, 5 mM **2a** in a solution of 20% (w/w) 4-methylpiperidine/DMF, was incubated at 37 °C for 30 minutes. Samples of the reaction mixtures were taken after 30 minutes and examined by LC-MS.

HPLC analysis was carried out on an Eclipse Plus C8 analytical column with Phase A/Phase B gradients [Phase A: MeOH with 0.1% trifluoroacetic acid, Phase B: $H_2O$ with 0.1% trifluoroacetic acid]. 50% Phase A in Phase B, 1 minute, and 50%–70% Phase A in Phase B, 4 minutes, then 70%–90% Phase A in Phase B, 3 minutes. The absorbance was monitored at 350 nm.

As shown in Fig. 5, 40% of tetrazine **2a** degrades in the 4-methylpiperidine solution after 30 minutes. In contrast, photocaged dihydrotetrazine **1a** is stable in the 4-methylpiperidine solution, and no detectable degradation was observed after 30 minutes.

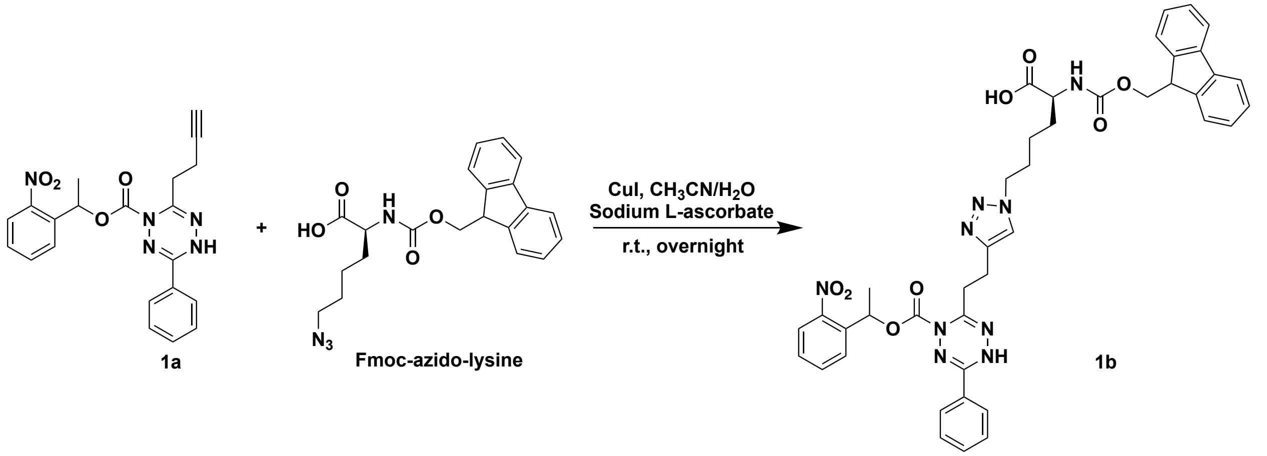


**Fig. 6. Synthesis of photocaged dihydrotetrazine-Fmoc-lysine 1b.**

Under argon, 7 mL of $CH_3CN/H_2O$ (v/v = 2/1) was added to a mixture of photocaged dihydrotetrazine **1a** (20 mg, 49 µmol), Fmoc-azido-lysine (39 mg, 100 µmol), CuI (35 mg, 184 µmol), and sodium L-ascorbate (14 mg, 71 µmol) at room temperature. The reaction mixture was stirred at room temperature overnight. Upon completion, the reaction solvent was removed under reduced pressure and the residue was dried under high vacuum. Then $CH_2Cl_2$ was added followed by filtration. The filtrate was concentrated by reduced pressure and the residue was purified by preparative thin layer chromatography on silica gel using 2%–20% $MeOH/CH_2Cl_2$ as the eluents, yielding the title compound **1b** as a pale-yellow solid (12 mg, 30%).

**(2S)-2-((((9H-fluoren-9-yl)methoxy)carbonyl)amino)-6-(4-(2-(4-((1-(2-nitrophenyl)ethoxy)carbonyl)-6-phenyl-1,4-dihydro-1,2,4,5-tetrazin-3-yl)ethyl)-1H-1,2,3-triazol-1-yl)hexanoic acid (1b)**

**$^1$H NMR** (500 MHz, $CD_2Cl_2$): δ 8.72 (s, 1 H), 7.94–7.24 (m, 17 H), 6.41–6.39 (m, 1 H), 5.81 (s, 1 H), 4.32–4.14 (m, 5 H), 3.66–3.54 (m, 1 H), 3.41 (s, 1 H), 3.14–2.86 (m,4 H), 1.74 (s, 3 H), 1.64 (s, 2 H), 1.26–0.88 (m, 4 H).

**$^{13}$C NMR** (126 MHz, $CD_2Cl_2$): δ 156.44, 155.99, 152.49, 150.47, 147.93, 144.61, 144.43, 144.20, 141.55, 141.52, 138.06, 134.28, 132.14, 129.21, 128.99, 128.86, 127.97, 127.79, 127.37, 127.02, 125.44, 124.77, 122.34, 120.28, 120.22, 70.94, 70.83, 67.06, 50.76, 49.72, 47.48, 31.31, 30.06, 29.38, 22.18, 21.35, 14.28.

**HRMS** *m/z* (ESI): calcd. for $C_{42}H_{42}N_9O_8$ $[M+H]^+$: 800.3151, found: 800.3141.

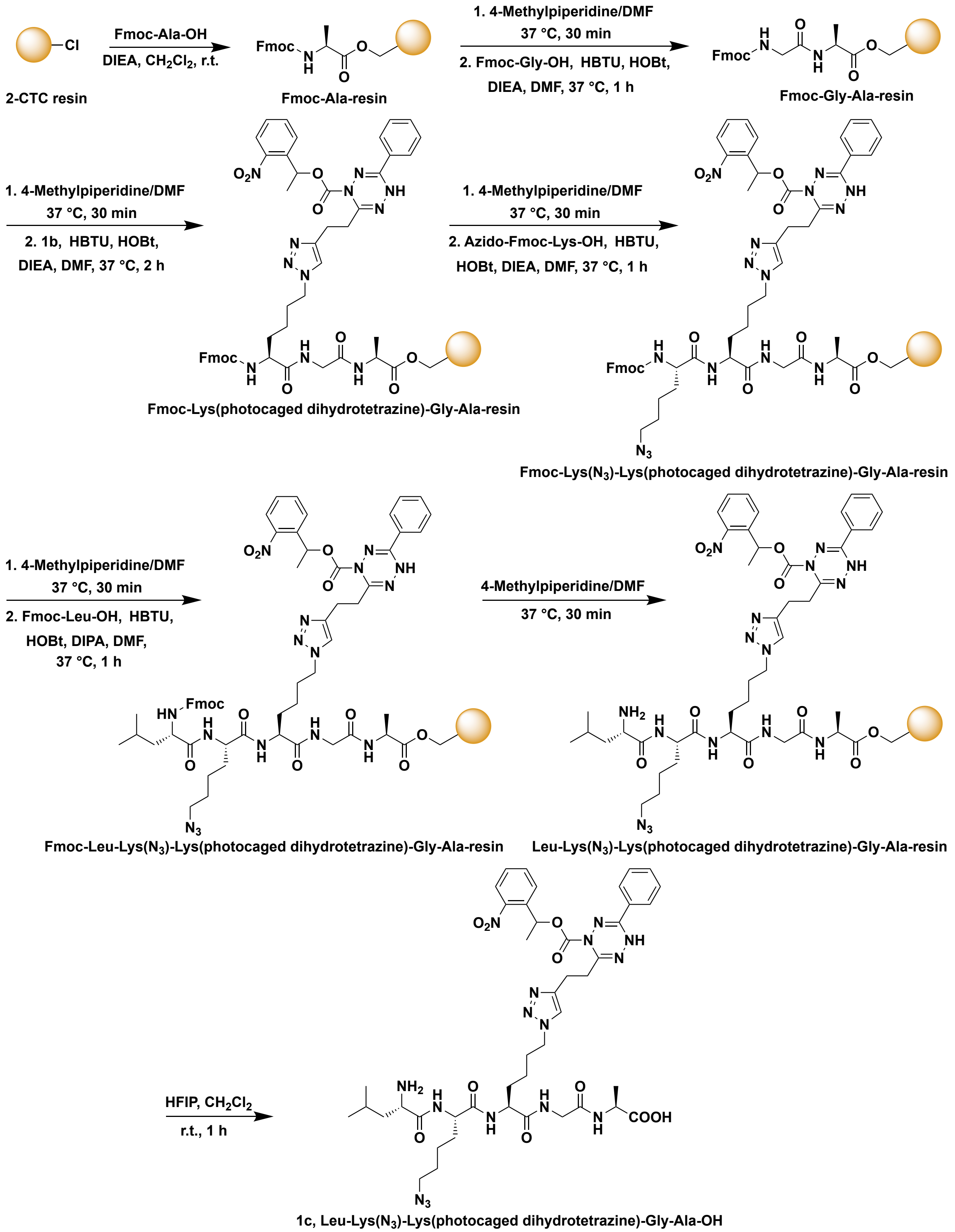


**Fig. 7. Synthesis of photocaged dihydrotetrazine-peptide 1c through solid-phase peptide synthesis (SPPS).[5]**

Photocaged dihydrotetrazine-peptide **1c** was prepared manually by following a standard Fmoc chemistry solid-phase peptide synthesis (SPPS) protocol. 2-Chlorotrityl chloride (2-CTC) resin (350 mg; loading: 1.14 mmol/g) was soaked in anhydrous $CH_2Cl_2$ (4 mL) for 30 minutes. The solvent was filtered off, and a solution of Fmoc-Ala-OH (373 mg, 1.2 mmol) and *N*,*N*-diisopropylethylamine (DIEA) (522 μL, 3 mmol) in anhydrous $CH_2Cl_2$ (4 mL) was added to the resin, which was shaken for 4 hours at room temperature. Then the solvent was filtered off and the resin was washed with $CH_2Cl_2$ (4 mL). A mixture of $CH_2Cl_2$/MeOH/DIEA (v/v/v = 8.5/1/0.5, 4 mL) was added and the resin was shaken for 30 minutes at room temperature, then washed with $CH_2Cl_2$ (3 × 4 mL) and $Et_2O$ (4 mL). The resin was dried under high vacuum for 5 hours, and the loading was determined by quantifying the absorbance of the Fmoc group. A small portion of the resin (2.67 mg) was treated with a solution of 20% (w/w) 4-methylpiperidine/DMF (1 mL) for 30 minutes at room temperature. To an aliquot of this solution (100 μL) was added 900 μL of DMF and the absorbance was read at 301 nm. The concentration of the dibenzofulvene-piperidine adduct was obtained by using the reported extinction coefficient (ε).[6] Thus, the resin loading was estimated to be 0.79 mmol/g. A portion of Fmoc-Ala-resin (115 mg, loading: 0.79 mmol/g) was used for the synthesis of the desired peptide. The Fmoc group was removed by treatment with a solution of 20% (w/w) 4-methylpiperidine/DMF (3 mL) at 37 °C for 30 minutes. The resin was washed with DMF (6 mL) and then treated with a solution of Fmoc-Gly-OH (119 mg, 0.4 mmol), HBTU (152 mg, 0.4 mmol), HOBt (69 mg, 0.4 mmol) and DIEA (139 μL, 0.8 mmol) in DMF (3 mL). The resin was shaken at 37 °C for 1 hour and then washed with DMF (3 mL), $CH_2Cl_2$ (3 × 4 mL) and $Et_2O$ (4 mL). The resin was dried under high vacuum overnight and 105 mg of Fmoc-Gly-Ala-resin was obtained. A portion of Fmoc-Gly-Ala-resin (18 mg) was used for the synthesis of the desired peptide. The Fmoc group was removed by treatment with a solution of 20% (w/w) 4-methylpiperidine/DMF (3 mL) at 37 °C for 30 minutes. The resin was washed with DMF (6 mL) and then treated with a solution of photocaged dihydrotetrazine-Fmoc-Lys-OH **1b** (12 mg, 15 μmol), HBTU (6 mg, 15 μmol), HOBt (2 mg, 15 μmol) and DIEA (5.2 μL, 30 μmol) in DMF (300 μL). The resin was shaken at 37 °C for 2 hours and then washed with DMF (3 mL), and $CH_2Cl_2$ (4 mL). The Fmoc group was then removed by treatment with a solution of 20% (w/w) 4-methylpiperidine/DMF (3 mL) at 37 °C for 30 minutes. Next, the resin was washed with DMF (6 mL) and then treated with a solution of azido-Fmoc-Lys-OH (18 mg, 45 μmol), HBTU (18 mg, 45 μmol), HOBt (6 mg, 45 μmol) and DIEA (15 μL, 900 μmol) in DMF (300 μL). The resin was shaken at 37 °C for 1 hour and then washed with DMF (3 mL), and $CH_2Cl_2$ (4 mL). Then, the Fmoc group was removed by treatment with a solution of 20% (w/w) 4-methylpiperidine/DMF (3 mL) at 37 °C for 30 minutes. Next, the resin was washed with DMF (6 mL) and then treated with a solution of Fmoc-Leu-OH (18 mg, 45 μmol), HBTU (18 mg, 45 μmol), HOBt (6 mg, 45 μmol) and DIEA (15 μL, 900 μmol) in DMF (300 μL). The resin was shaken at 37 °C for 1 hour and then washed with DMF (3 mL), and $CH_2Cl_2$ (4 mL). Then, the Fmoc group was removed by treatment with a solution of 20% (w/w) 4-methylpiperidine/DMF (3 mL) at 37 °C for 30 minutes. Next, the protected peptide was released from the resin by treatment with a freshly prepared solution of 20% (v/v) 1,1,1,3,3,3-hexafluoro-2-propanol (HFIP)/ $CH_2Cl_2$ (1 mL) at room temperature for 1 hour followed by filtration. The resin was washed with $CH_2Cl_2$ (2 × 1 mL) and the combined fractions were evaporated in vacuo and dried under high vacuum overnight affording Leu-Lys($N_3$)-Lys(photocaged dihydrotetrazine)-Gly-Ala-OH **1c** as a pale-yellow solid (5.1 mg, 50%). **HRMS** of **1c** *m/z* (ESI): calcd. for $C_{44}H_{61}N_{16}O_{10}$ $[M+H]^+$: 973.4751, found: 973.4749.

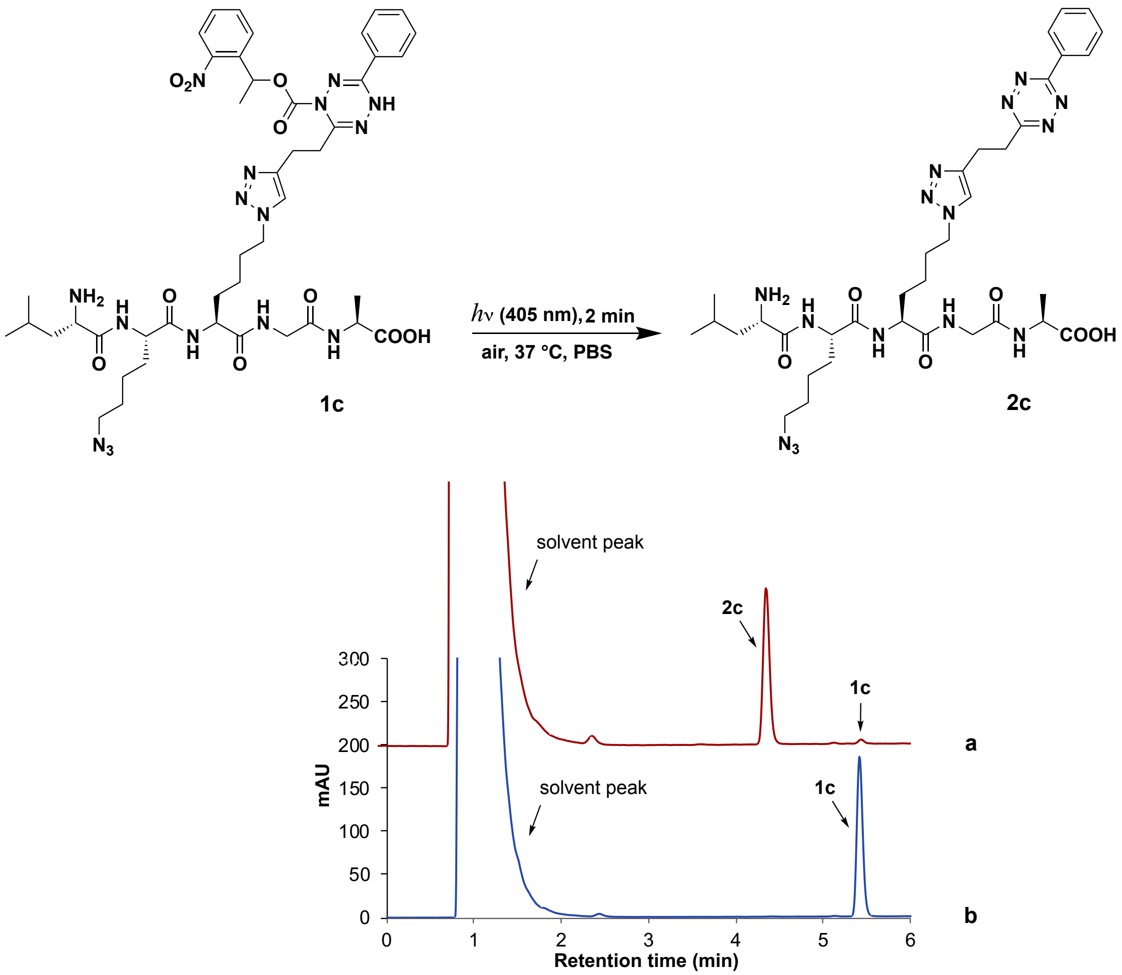


**Fig. 8. Photo-activation of tetrazine on peptide.** The reaction was carried out by irradiation of photocaged dihydrotetrazine-peptide **1c** (10 μM) with LED light (405 nm, 18 W) in PBS solution (containing 0.2% DMSO and 0.2% DMF) open to air at 37 °C for 2 minutes. Samples were taken from the reaction mixture and examined by LC-MS. **a**, After irradiation by LED light for 2 minutes (red spectrum). **b**, Before irradiation by LED light (blue spectrum).

HPLC/ELSD analysis was carried out on an Eclipse Plus C8 analytical column with Phase A/Phase B gradients [Phase A: MeOH with 0.1% trifluoroacetic acid, Phase B: $H_2O$ with 0.1% trifluoroacetic acid]. 50% Phase A in Phase B, 1 minute, and 50%–80% Phase A in Phase B, 4 minutes, then 80%–90% Phase A in Phase B, 3 minutes. ELSD spectra of samples at were presented.

As shown in Fig. 8, Leu-Lys($N_3$)-Lys(tetrazine)-Gly-Ala-OH **2c** was generated from the dihydrotetrazine-peptide **1c** after 2 minutes of irradiation with a 96% yield. **HRMS** of **2c** *m/z* (ESI): calcd. for $C_{35}H_{52}N_{15}O_6$ $[M+H]^+$: 778.4220, found: 778.4234.

#### 5. Spatiotemporal Live-Cell Labeling with Alexa Fluor Dyes

##### 5.1. Synthesis of photocaged dihydrotetrazine-diacylphospholipid 1d

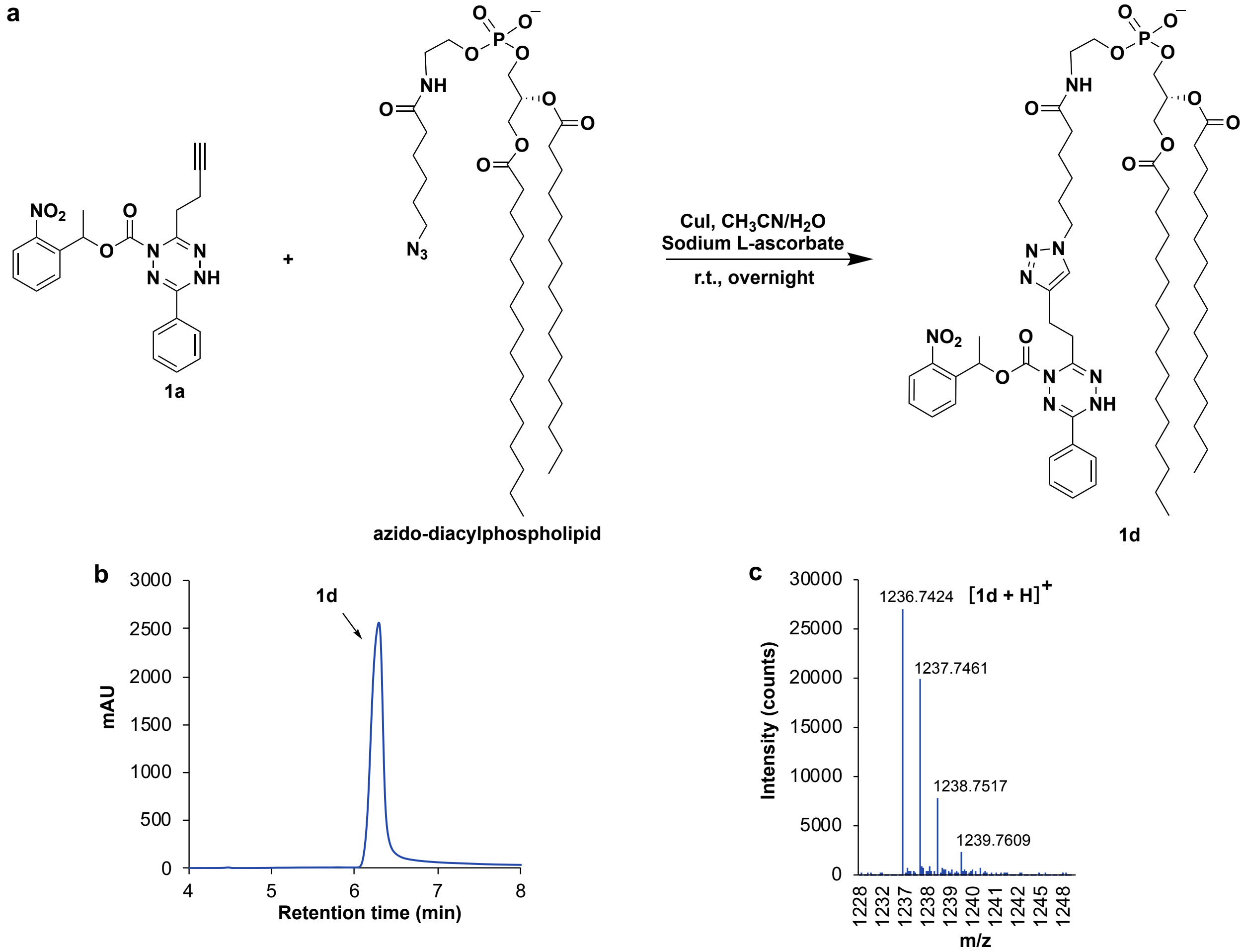


**Fig. 9. Synthesis of photocaged dihydrotetrazine-diacylphospholipid 1d. a,** Under argon, 1.2 mL of $CH_3CN/H_2O$ (v/v = 2/1) was added to a mixture of photocaged dihydrotetrazine **1a** (7.1 mg, 18 μmol), azido-diacylphospholipid (20 mg, 24 μmol), CuI (6 mg, 32 μmol), and sodium L-ascorbate (2.4 mg, 12 μmol) at room temperature. The reaction mixture was stirred at room temperature overnight. Upon completion, the reaction solvent was removed under reduced pressure and the residue was dried over high vacuum. Then $CH_2Cl_2$ was added, followed by filtration. The filtrate was concentrated by reduced pressure and the residue was purified by preparative thin layer chromatography on silica gel using 10% $MeOH/CH_2Cl_2$ as the eluent yielding the title compound **1d** as a pale-yellow solid (11.5 mg, 53%), which was examined by LC-MS. **b,** HPLC/ELSD spectrometry of **1d**. **c,** HRMS spectrometry of **1d**. **HRMS** of **1d** *m/z* (ESI): calcd. for $C_{64}H_{103}N_9O_{13}P$ $[M+H]^+$: 1236.7407, found: 1236.7424.

HPLC/ELSD analysis was carried out on an Eclipse Plus C8 analytical column with Phase A/Phase B gradients [Phase A: MeOH with 0.1% trifluoroacetic acid, Phase B: $H_2O$ with 0.1% trifluoroacetic acid]. 50% Phase A in Phase B, 1 minute, and 50%–95% Phase A in Phase B, 2 minutes, 95%–100% Phase A in Phase B, 1 minute, 100% Phase A, 4 minutes.

#### 5.2. Light-activated ligation between photocaged dihydrotetrazine-diacylphospholipid 1d and fluorophore TCO-AF488 3a

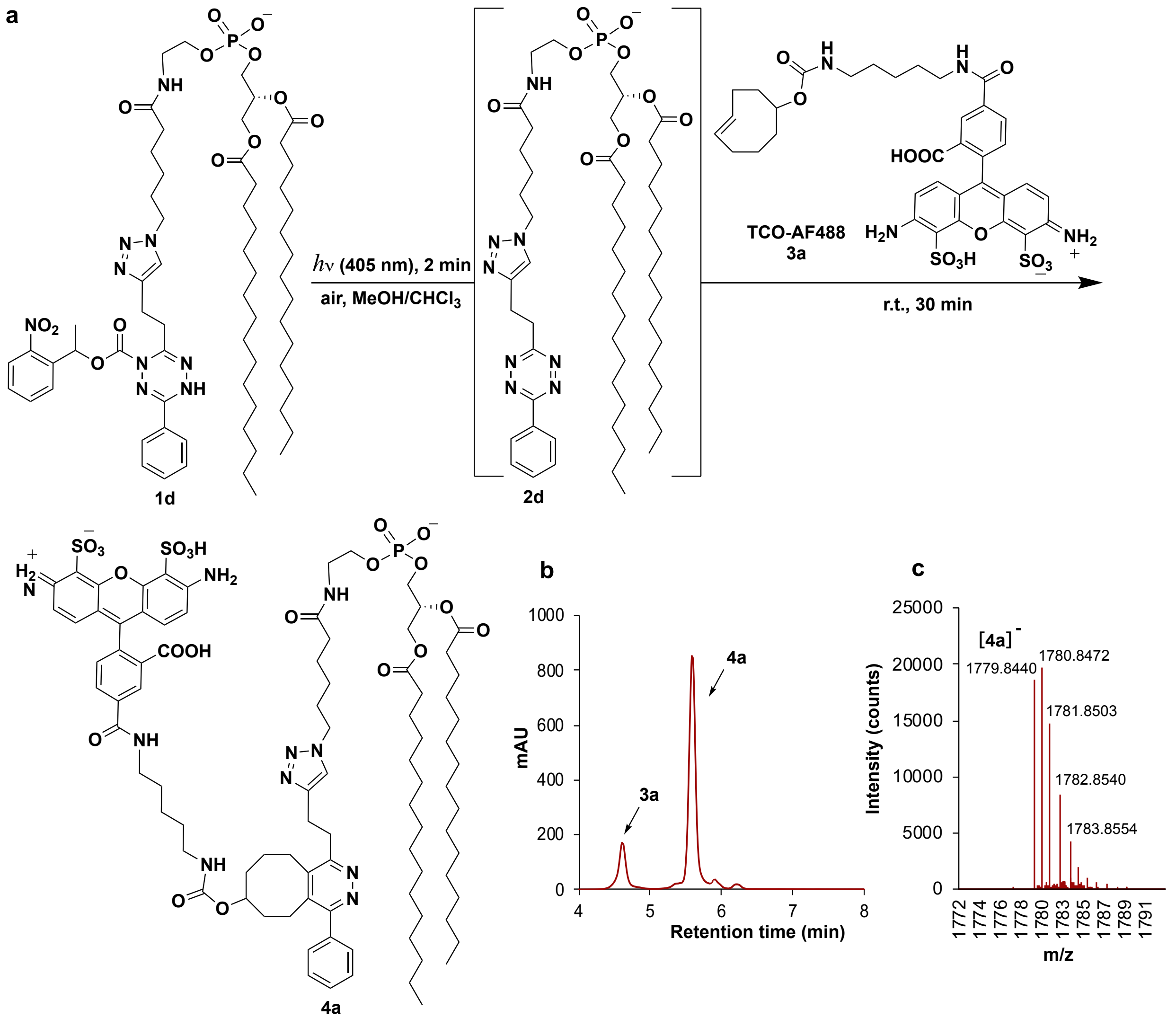


**Fig. 10. Light-triggered tetrazine ligation between photocaged dihydrotetrazine-diacylphospholipid 1d and *trans*-cyclooctene carbamate caged Alexa Fluor 488 dye 3a. a,** The reaction was started by irradiating a solution of photocaged dihydrotetrazine-diacylphospholipid **1d** (0.38 µmol) and *trans*-cyclooctene carbamate caged Alexa Fluor 488 dye (TCO-AF488) **3a** (0.26 µmol) in $MeOH/CHCl_3$ (200 µL/20 µL) using LED light (405 nm, 18 W) in an open vial for 2 minutes at room temperature. After irradiation, the reaction mixture was incubated for 30 minutes. Purification of the conjugated product **4a** was performed by preparative thin layer chromatography on silica gel using 20% $MeOH/CH_2Cl_2$ as the eluent. LC-MS analysis was carried out. **b,** HPLC/ELSD spectrometry of **4a**. **c,** HRMS spectrometry of **4a**. **HRMS** of **4a** *m/z* (ESI): calcd. for $C_{90}H_{128}N_{10}O_{21}PS_2$ $[M]^-$: 1779.8440, found: 1779.8440.

HPLC/ELSD analysis was carried out on an Eclipse Plus C8 analytical column with Phase A/Phase B gradients [Phase A: MeOH with 0.1% trifluoroacetic acid, Phase B: $H_2O$ with 0.1%

trifluoroacetic acid]. 50% Phase A in Phase B, 1 minute, and 50%–95% Phase A in Phase B, 2 minutes, 95%–100% Phase A in Phase B, 1 minute, 100% Phase A, 4 minutes.

##### 5.3. Spatiotemporal labeling of cell membranes by light-activated tetrazine ligations

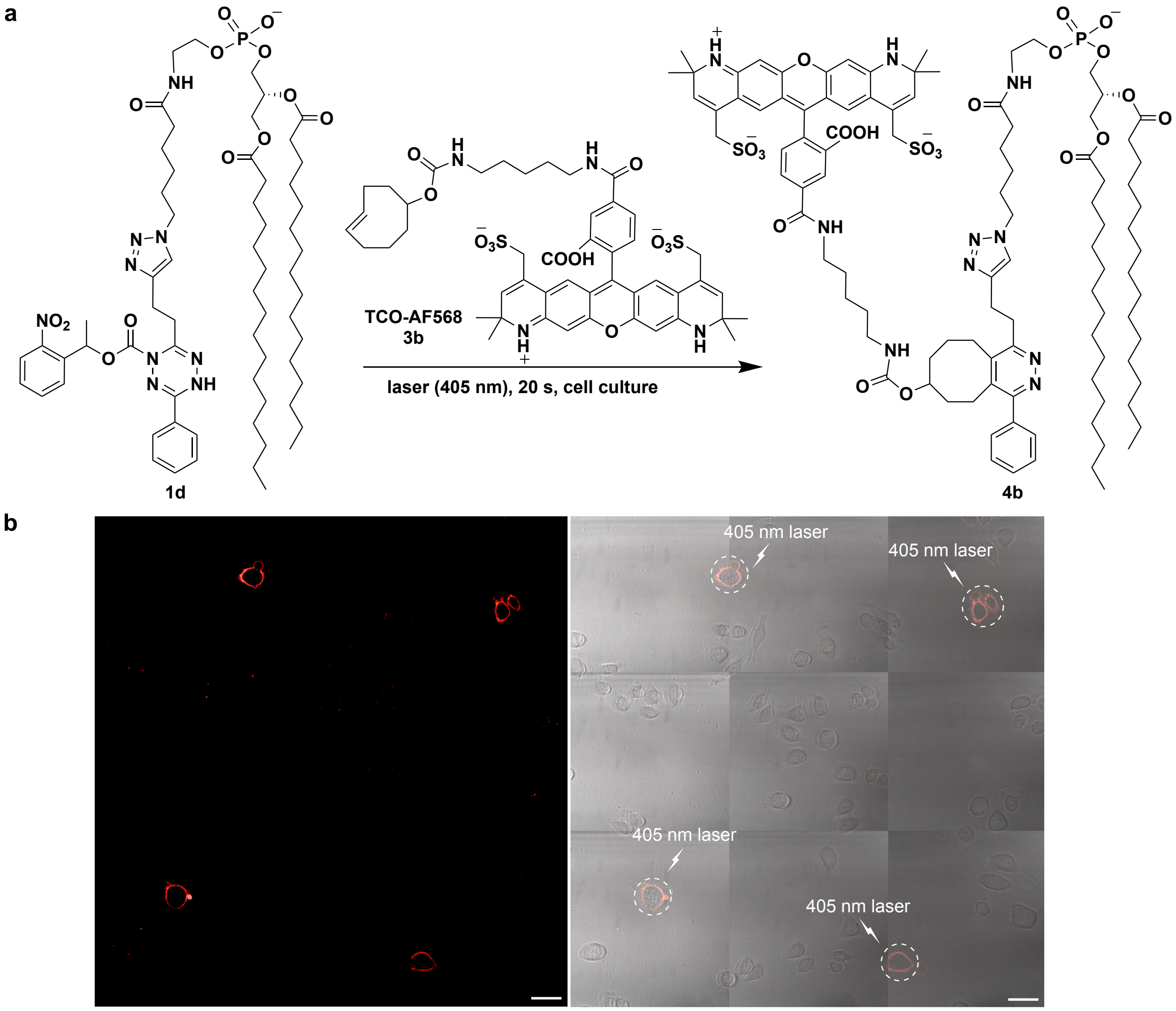


**Fig. 11. Spatiotemporal cell membrane labeling of living HeLa S3 cancer cells by light-activated tetrazine ligations. a,** Light-activated ligations between photocaged dihydrotetrazine-diacylphospholipid **1d** and *trans*-cyclooctene carbamate caged Alexa Fluor 568 dye (TCO-AF568) **3b** in the presence of living HeLa S3 cancer cells. **b,** 3*3 tiled (total area = 0.75 mm by 0.75 mm) fluorescence imaging of cell membranes of living HeLa S3 cells using Alexa Fluor 568 dye (AF568) after light-activated tetrazine ligations between photocaged dihydrotetrazine-diacylphospholipid **1d** and TCO-AF568 **3b**. Left image: Fluorescence channel; Right image: Merged fluorescence and brightfield channels. The white dashed lines in the merged image denote the 405 nm laser irradiated regions. Scale bar: 50 μm.

Human HeLa S3 cancer cells (ATCC, CCL-2.2) were grown in the complete media of Dulbecco's Modified Eagle's Minimal Essential Medium (DMEM) with high glucose (Life Technologies—Gibco, 11995073) supplemented with 10% heat-inactivated fetal bovine serum (Omega Scientific, FB02), 100 U/mL penicillin, and 100 μg/mL streptomycin (Life

Technologies—Gibco, 15140122). Cells were grown at 37 °C in a humidified atmosphere containing 5% $CO_2$. An 8-well chamber slide (cat#80826, ibidi USA Inc., Fitchburg, Wisconsin) was pre-coated with 0.5% (w/v) poly-lysine in $H_2O$ to facilitate cell adhesion during imaging. HeLa S3 cell lines were plated at 40% density per well in 200 μL of DMEM complete media. 24 hours later, the cells were treated as follow: HeLa S3 cancer cells were incubated in 200 μL of photocaged dihydrotetrazine-diacylphospholipid **1d** (60 nM) in PBS solution (containing 0.1% DMSO) at 37 °C for 5 minutes. Excess **1d** was then removed by washing the cells with fresh PBS solution. Next, the washed cells were incubated with 3 nM TCO-AF568 **3b** in PBS solution (containing 0.1% DMSO). To trigger the activation of the photocaged dihydrotetrazine and the subsequent bioorthogonal tetrazine ligation, selected cells were laser irradiated (405 nm, 20 mW) for 20 seconds using a ZEISS 880 laser scanning microscope. Five minutes after the laser uncaging event, cells were washed with fresh full-growth media and were imaged by fluorescence microscopy.

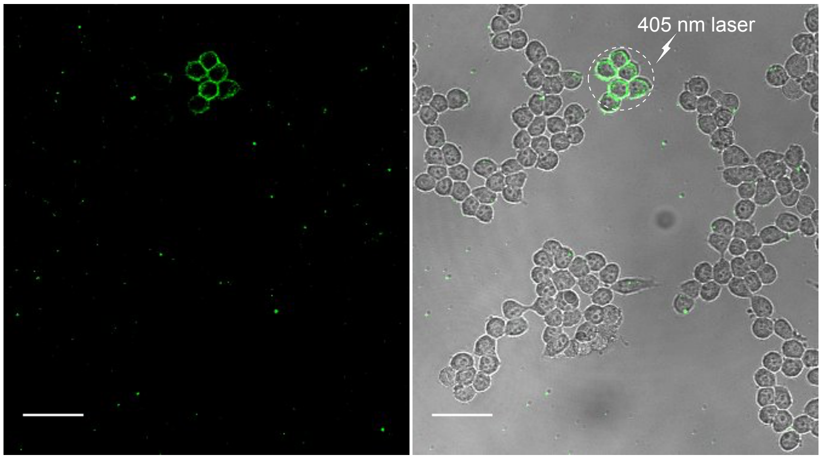


**Fig. 12. Spatiotemporal labeling of living Hep 3B cancer cell membranes by light-activated tetrazine ligations.** Fluorescence imaging of cell membranes of living Hep 3B cancer cells using Alexa Fluor 488 dye after light-activated tetrazine ligations between photocaged dihydrotetrazine-diacylphospholipid **1d** and TCO-AF488 **3a**. Left image: Fluorescence channel; Right image: Merged fluorescence and brightfield channels. The white dashed lines in the merged image denote the 405 nm laser irradiated regions. Scale bar: 50 μm.

Human Hep 3B hepatocellular carcinoma cells (ATCC, HB-8064) were grown in the complete media of Eagle's Minimal Essential Medium (EMEM) with high glucose (Life Technologies—Gibco, 11995073) supplemented with 10% heat-inactivated fetal bovine serum (Omega Scientific, FB02), 100 U/mL penicillin, and 100 μg/mL streptomycin (Life Technologies—Gibco, 15140122). Cells were grown at 37 °C in a humidified atmosphere containing 5% $CO_2$. An 8-well chamber slide (cat#80826, ibidi USA Inc., Fitchburg, Wisconsin) was pre-coated with 0.5% (w/v) poly-lysine in $H_2O$ in order to limit cell movement during imaging. Hep 3B cells were plated at 30,000 cells per well on 8-well plates in 200 μL of EMEM complete media and incubated for 24 hours to promote cell adhesion and growth. Next, cells were incubated in 200 μL of photocaged dihydrotetrazine-diacylphospholipid **1d** (60 nM) in PBS solution (containing 0.1% DMSO) at 37 °C for 5 minutes. Excess of **1d** was then removed by washing the cells with PBS solution. Then, the washed cells were incubated with 3 nM of TCO-AF488 **3a** in PBS solution (containing 0.1% DMSO). To trigger the activation of photocaged dihydrotetrazine-diacylphospholipid **1d** and the subsequent bioorthogonal tetrazine ligation with TCO-AF488 **3a**, selected cells were laser irradiated (405 nm, 20 mW) for 20 seconds using a laser scanning microscope. Five minutes after the laser uncaging event, cells were washed with complete media and were imaged by fluorescence microscopy.

#### 6. Delivery of Doxorubicin by Light-Activated Tetrazine Ligation

##### 6.1. Synthesis of *trans*-cyclooctene carbamate caged doxorubicin (TCO-Dox) 3c

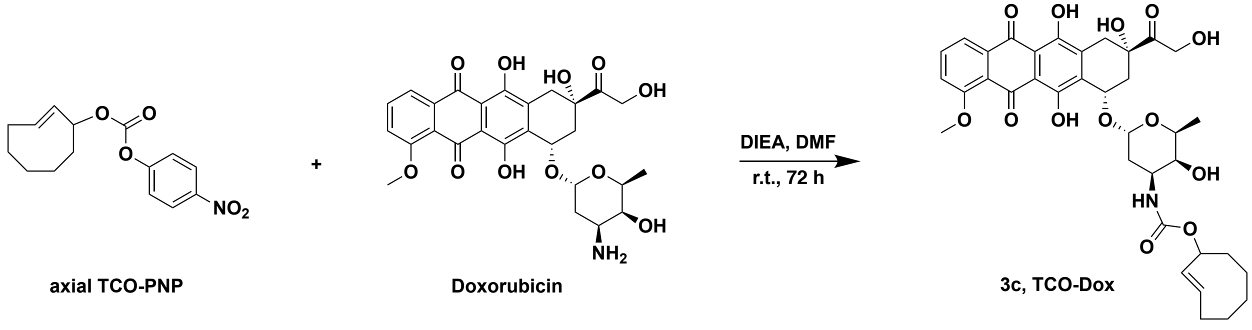


**Fig. 13. Synthesis of *trans*-cyclooctene carbamate caged doxorubicin (TCO-Dox) 3c.**
DIEA (0.22 mL, 1.22 mmol) was added dropwise to a mixture of axial TCO-PNP (40 mg, 0.137 mmol) and doxorubicin hydrochloride (90 mg, 0.155 mmol) in anhydrous DMF (6.3 mL). The reaction mixture was stirred at room temperature for 72 hours. Upon completion, the reaction mixture was concentrated under reduced pressure. The residue was purified by column chromatography on silica gel using 0.5%–2% MeOH/$CH_2Cl_2$ as the eluent yielding the title compound **3c** as a red solid (24 mg, 25%).

***Trans*-cyclooctene carbamate caged doxorubicin (3c)**
**$^1$H NMR** (500 MHz, $CD_3OD$): δ 13.95 (s, 1 H), 13.20 (s, 1 H), 8.01 (t, *J* = 5 Hz, 1 H), 7.78 (dd, *J* = 10.0 Hz, *J* = 5.0 Hz, 1 H), 7.38 (dd, *J* = 10.0 Hz, *J* = 5.0 Hz, 1 H), 5.74 (m, 1 H), 5.50 (m, 2 H), 5.27 (m, 2 H), 5.13 (m, 1 H), 4.75 (m, 2 H), 4.53 (brs, 1 H), 4.10 (s, 4 H), 3.86 (s, 1 H), 3.68 (s, 1 H), 3.23 (d, *J* = 19 Hz, 1 H), 2.95 (d, *J* = 19 Hz, 2 H), 2.42 (brs, 1 H), 2.31 (d, *J* = 15 Hz, 1 H), 2.16 (m, 1 H), 2.03-1.75 (m, 6 H), 1.61 (m, 2 H), 1.44 (m, 1 H), 1.29 (m, 3 H), 1.02 (m, 1 H), 0.74 (m, 1 H)
**$^{13}$C NMR** (126 MHz, $CD_3OD$): δ 214.10, 187.17, 186.74, 161.11, 156.29, 155.74, 155.15, 135.91, 135.55, 133.70, 133.65, 131.81, 131.34, 120.88, 119.95, 118.52, 111.65, 111.46, 100.85, 74.23, 69.79, 69.70, 67.42, 65.70, 56.79, 53.59, 46.89, 40.70, 36.03, 35.98, 35.75, 34.08, 30.31, 29.83, 29.15, 24.15, 16.99.
**HRMS** *m/z* (ESI): calcd. for $C_{36}H_{41}N_1O_{13}Na_1$ $[M+Na]^+$: 718.2471; found: 718.2470.

##### 6.2. Doxorubicin release by light-activated tetrazine ligation in PBS solution

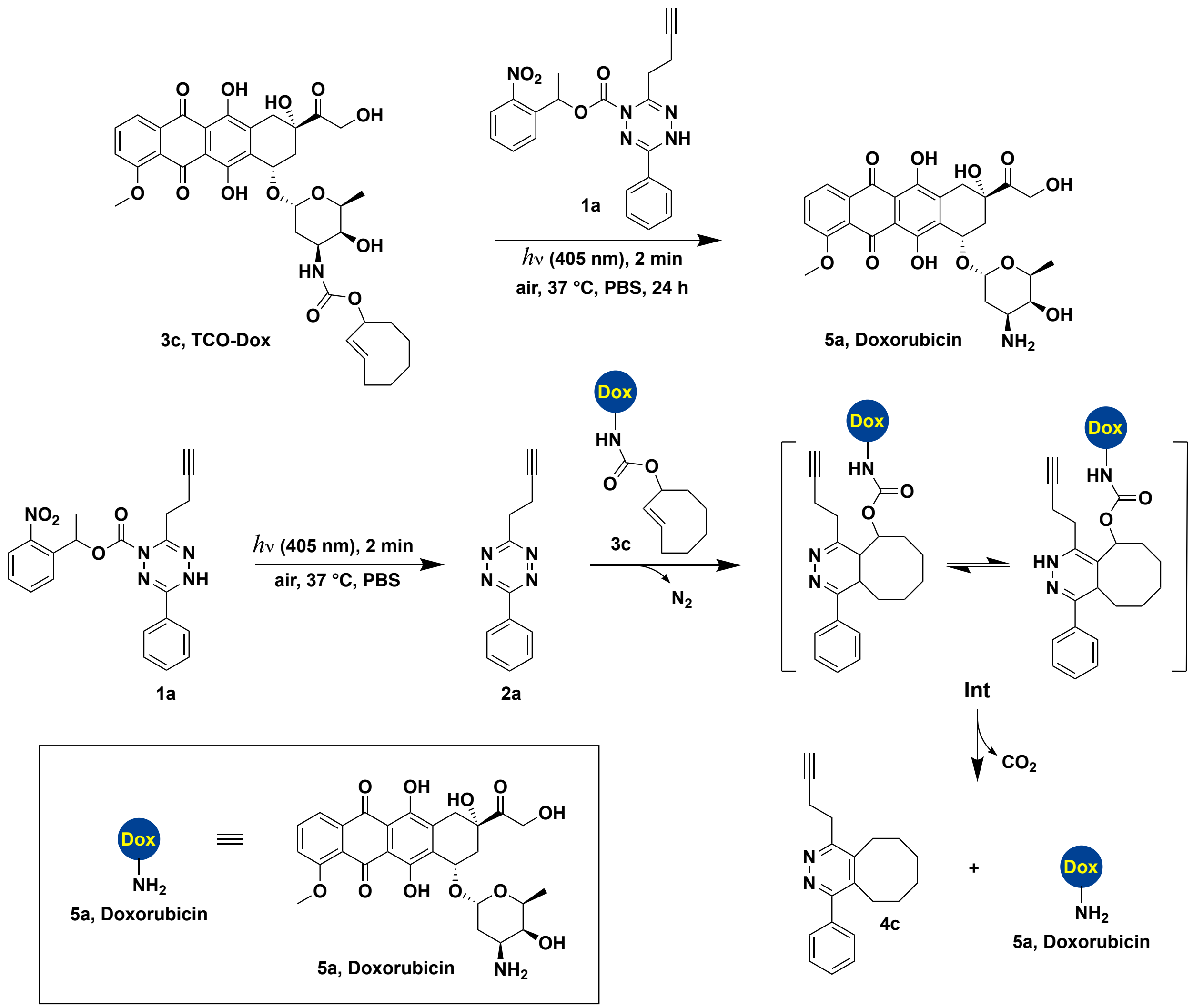


**Fig. 14. Light-activated tetrazine ligation with TCO-doxorubicin 3c.** The reaction was initiated by irradiating a mixture of photocaged dihydrotetrazine **1a** (8 µM) and *trans*-cyclooctene carbamate caged doxorubicin **3c** (TCO-Dox) (5.5 µM) in PBS solution (containing 0.1% DMSO) with LED light (405 nm, 18W) for 2 minutes in an open vial. The reaction mixture was then incubated for 24 hours at 37 °C. Samples of the reaction mixture were taken at different time points and examined by LC-MS.

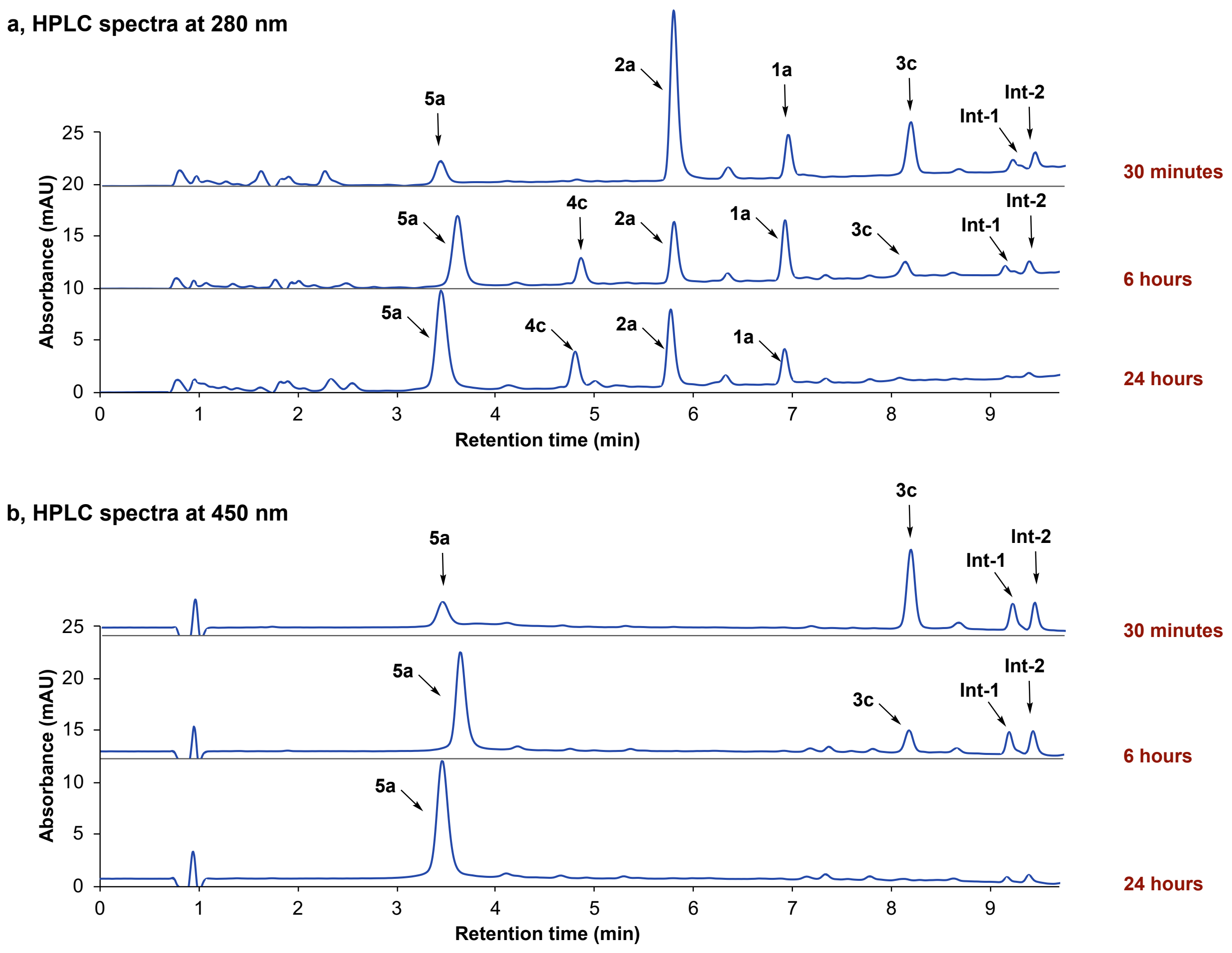


**Fig. 15. HPLC analysis of light-activated tetrazine ligation between photocaged dihydrotetrazine 1a and TCO-doxorubicin 3c. a,** HPLC spectra with absorbance monitored at 280 nm. **b,** HPLC spectra with absorbance monitored at 450 nm. The reaction was initiated by irradiating a solution of photocaged dihydrotetrazine **1a** (8 μM) and *trans*-cyclooctene carbamate caged doxorubicin **3c** (TCO-Dox) (5.5 μM) in PBS solution (containing 0.1% DMSO) by LED light (405 nm, 18W) for 2 minutes in an open vial at 37 °C. The reaction mixture was then incubated at 37 °C for 24 hours. Samples of the reaction mixture were taken at different time points (30 minutes, 6 hours, and 24 hours) and examined by LC-MS. **1a**: photocaged dihydrotetrazine; **2a**: tetrazine; **3c**: *trans*-cyclooctene carbamate caged doxorubicin; **4c**: pyridazine elimination product; **5a**: doxorubicin. **Int:** iEDDA-adduct and its tautomer. Two iEDDA-adduct peaks of **Int-1** and **Int-2**, were detected.

HPLC analysis was carried out on an Eclipse Plus C8 analytical column with Phase A/Phase B gradients [Phase A: MeOH with 0.1% trifluoroacetic acid, Phase B: $H_2O$ with 0.1% trifluoroacetic acid]. 50% Phase A in Phase B, 1 minute, and 50%–70% Phase A in Phase B, 4 minutes, 70%–90% Phase A in Phase B, 3 minutes, 90%–50% Phase A in Phase B, 0.5 minute, then 70%–90% Phase A in Phase B, 3 minutes.

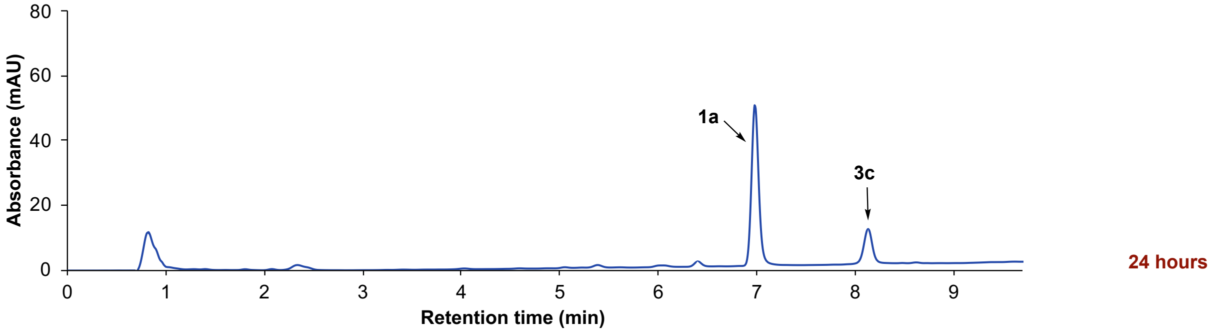


**Fig. 16. HPLC analysis of a mixture of photocaged dihydrotetrazine 1a and *trans*-cyclooctene carbamate caged doxorubicin 3c in the dark.** In the absence of light, a mixture of photocaged tetrazine **1a** (8 μM) and *trans*-cyclooctene carbamate caged doxorubicin **3c** (TCO-Dox) (5.5 μM) in PBS solution (containing 0.1% DMSO) was incubated for 24 hours in an open vial at 37 °C. A sample of the reaction mixture was taken after 24 hours and was examined by LC-MS.

LC-MS analysis was carried out on an Eclipse Plus C8 analytical column with Phase A/Phase B gradients [Phase A: MeOH with 0.1% trifluoroacetic acid, Phase B: $H_2O$ with 0.1% trifluoroacetic acid]. 50% Phase A in Phase B, 1 minute, and 50%–70% Phase A in Phase B, 4 minutes, 70%–90% Phase A in Phase B, 3 minutes, 90%–50% Phase A in Phase B, 0.5 minute, then 70%–90% Phase A in Phase B, 3 minutes. The absorbance was monitored at 280 nm.

##### 6.3. Doxorubicin delivery to cancer cells enabled by light-activated tetrazine ligation

The following solutions were prepared: PBS solution (containing 0.1% DMSO); complete media EMEM with high glucose (Life Technologies—Gibco, 11995073) supplemented with 10% heat-inactivated fetal bovine serum (Omega Scientific, FB02), 100 U/mL penicillin, and 100 µg/ml streptomycin (Life Technologies—Gibco, 15140122); photocaged tetrazine **1a** (16 µM) in PBS solution (containing 0.1% DMSO); *trans*-cyclooctene carbamate caged doxorubicin **3c** (11 µM) in EMEM complete media; doxorubicin **5a** (11 µM) in EMEM complete media.

Human Hep 3B hepatocellular carcinoma cells (ATCC, HB-8064) were grown in EMEM complete media. Cells were maintained at 37 °C in a humidified atmosphere containing 5% $CO_2$. Hep 3B cell lines were plated at 30,000 cells per well on 96-well plates in 200 of µL EMEM complete media. 24 hours later, the cells were treated as follows:

**Control**: Cell media was exchanged with 100 µL of PBS solution (containing 0.1% DMSO). Next, cells were irradiated by LED light (405 nm, 18W) for 2 minutes. Cells were subsequently incubated at 37 °C for 20 minutes, followed by the addition of 100 µL of EMEM complete media.

**No treatment**: Cell culturing media was exchanged with 200 µL of EMEM complete media.

**1a + 3d (No light)**: Cell culturing media was exchanged with 100 µL of **1a** (16 µM) in PBS solution (containing 0.1% DMSO). Cells were incubated at 37 °C for 20 minutes. Next, 100 µL of **3c** (11 µM) in EMEM complete media was added.

**1a + 3d (*h*ν 405 nm)**: Cell culturing media was exchanged with 100 µL of **1a** (16 µM) in PBS solution (containing 0.1% DMSO). Next, cells were irradiated with LED light (405 nm, 18W) for 2 minutes. Cells were subsequently incubated at 37 °C for 20 minutes, followed by the addition of 100 µL of EMEM complete media.

**5a (*h*ν 405 nm)**: Cell culturing media was exchanged with 100 µL of PBS solution (containing 0.1% DMSO). Next, cells were irradiated with LED light (405 nm, 18W) for 2 minutes. Cells were incubated at 37 °C for 20 minutes, followed by the addition of 100 µL of **5a** (11 µM) in EMEM complete media.

**1a (*h*ν 405 nm)**: Cell culturing media was exchanged with 100 µL of **1a** (16 µM) in PBS solution (containing 0.1% DMSO). Next, cells were irradiated with LED light (405 nm, 18W) for 2 minutes. Cells were subsequently incubated at 37 °C for 20 minutes, followed by the addition of 100 µL of EMEM complete media.

**3d (*h*ν 405 nm)**: Cell culturing media was exchanged with 100 µL of PBS solution (containing 0.1% DMSO). Next, cells were irradiated with LED light (405 nm, 18W) for 2 minutes. Cells were subsequently incubated at 37 °C for 20 minutes, followed by the addition of 100 µL of **3c** (11 µM) in EMEM complete media.

After the treatments described above, cells were incubated at 37 °C with 5% $CO_2$ for 24 hours. To quantify cell viability, a standard WST-1 Assay was performed (CELLPRO-RO, Millipore Sigma, SKU 11644807001). Following the manufacture's protocol, cell culturing media was exchanged with 100 µL of EMEM containing 10 µL of WST-1 Reagent. The cells were subsequently incubated at 37 °C for 1 hour, followed by absorbance measurement at 450 nm using a plate reader. This cell viability assay was performed in triplicate.

#### 6.4. Mass spectra of HPLC/HRMS analysis of the light-activated tetrazine ligation enabled doxorubicin release

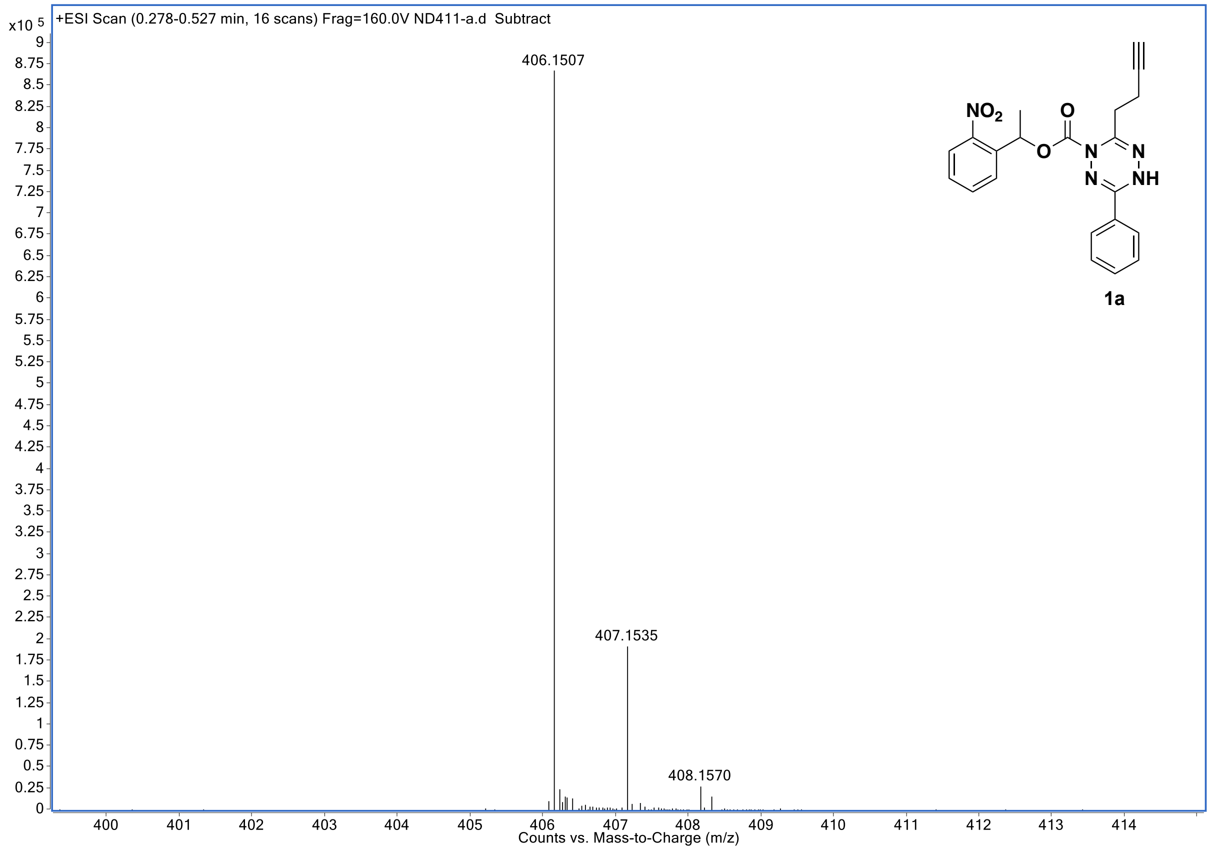

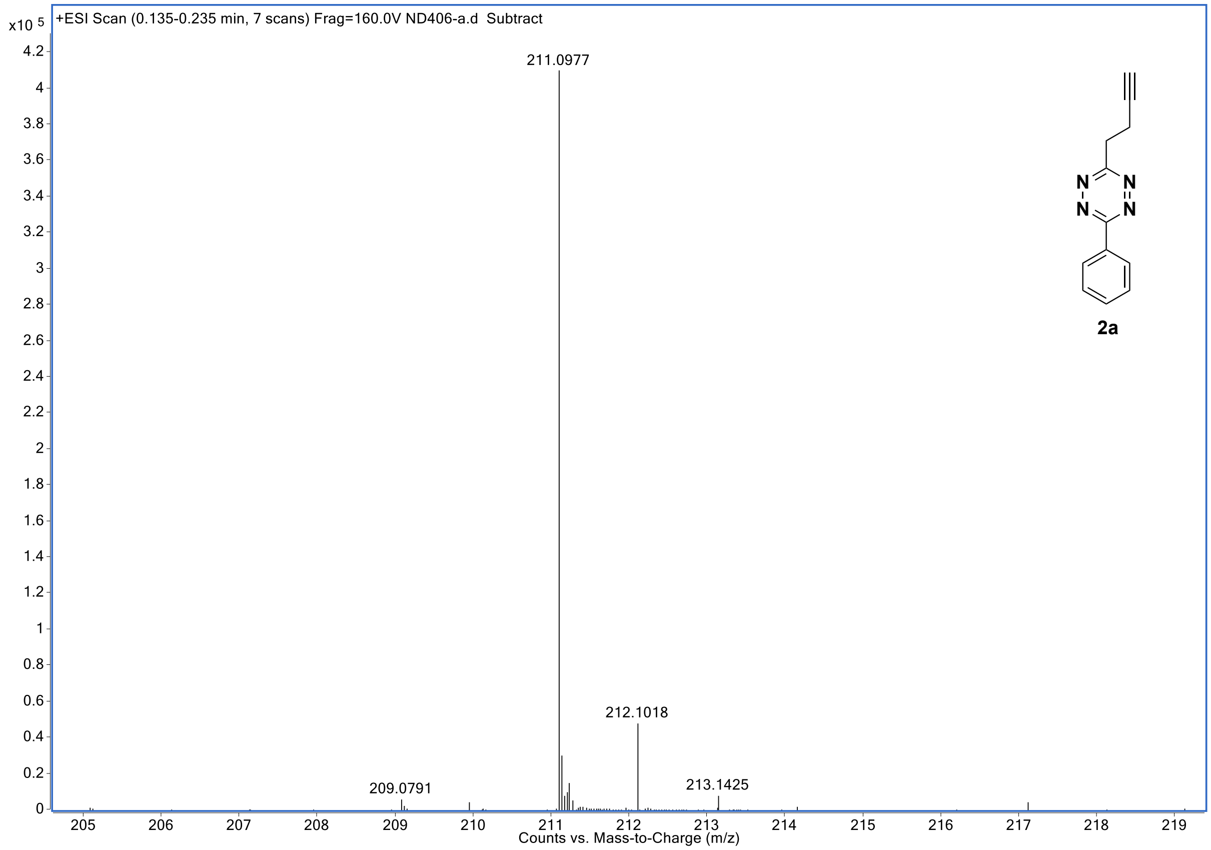
+ESI Scan (0.135-0.235 min, 7 scans) Frag=160.0V ND406-a.d Subtract
x10 5
211.0977
212.1018
209.0791
213.1425
Counts vs. Mass-to-Charge (m/z)
2a

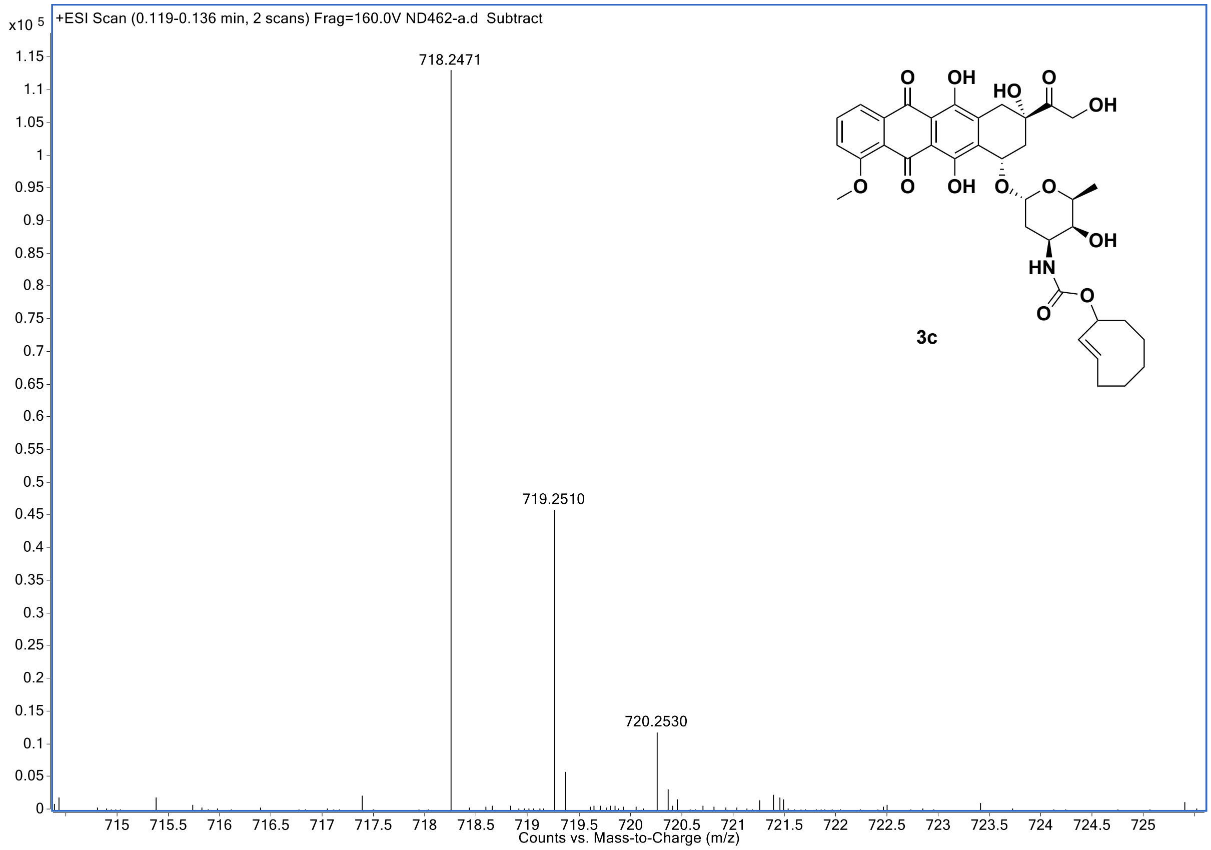
+ESI Scan (0.119-0.136 min, 2 scans) Frag=160.0V ND462-a.d Subtract
x10 5
718.2471
719.2510
720.2530
Counts vs. Mass-to-Charge (m/z)
3c

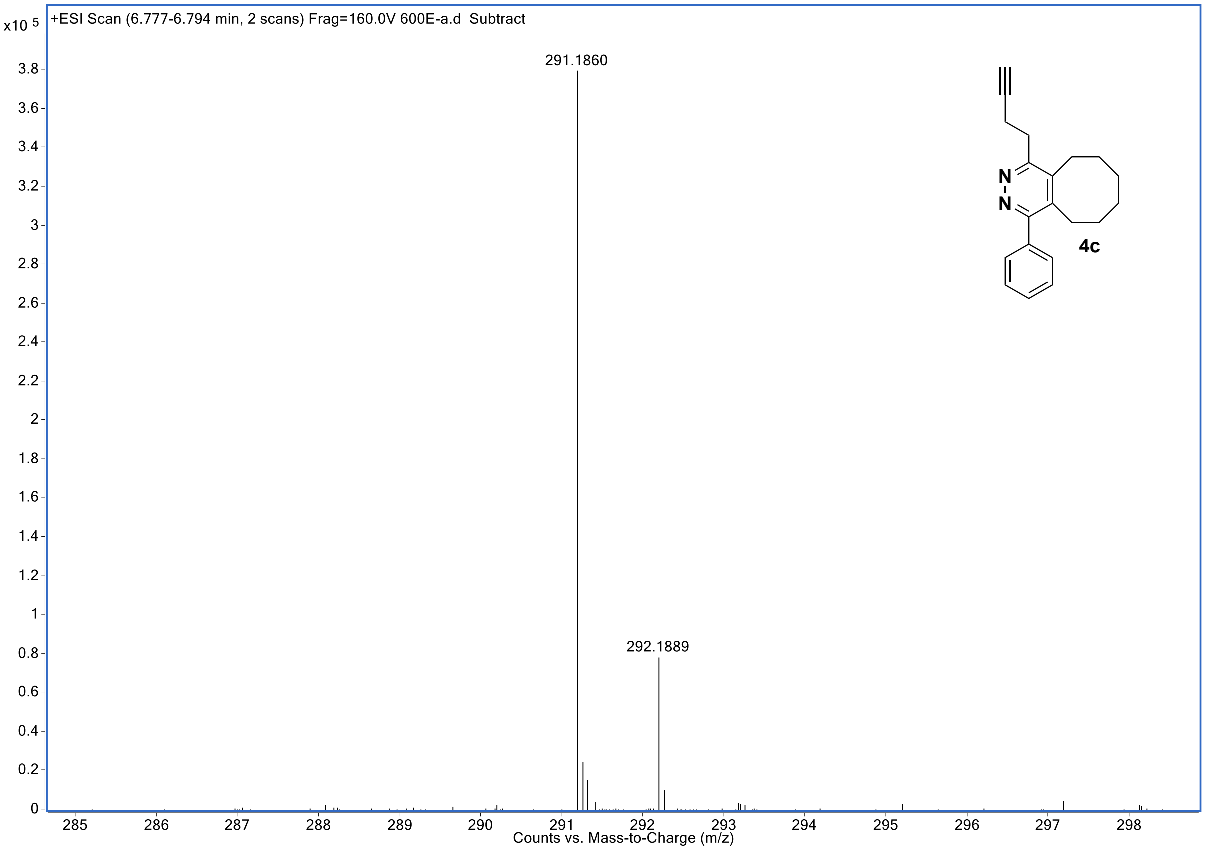

+ESI Scan (6.777-6.794 min, 2 scans) Frag=160.0V 600E-a.d Subtract
x10 5
291.1860
292.1889
N
N
4c
Counts vs. Mass-to-Charge (m/z)


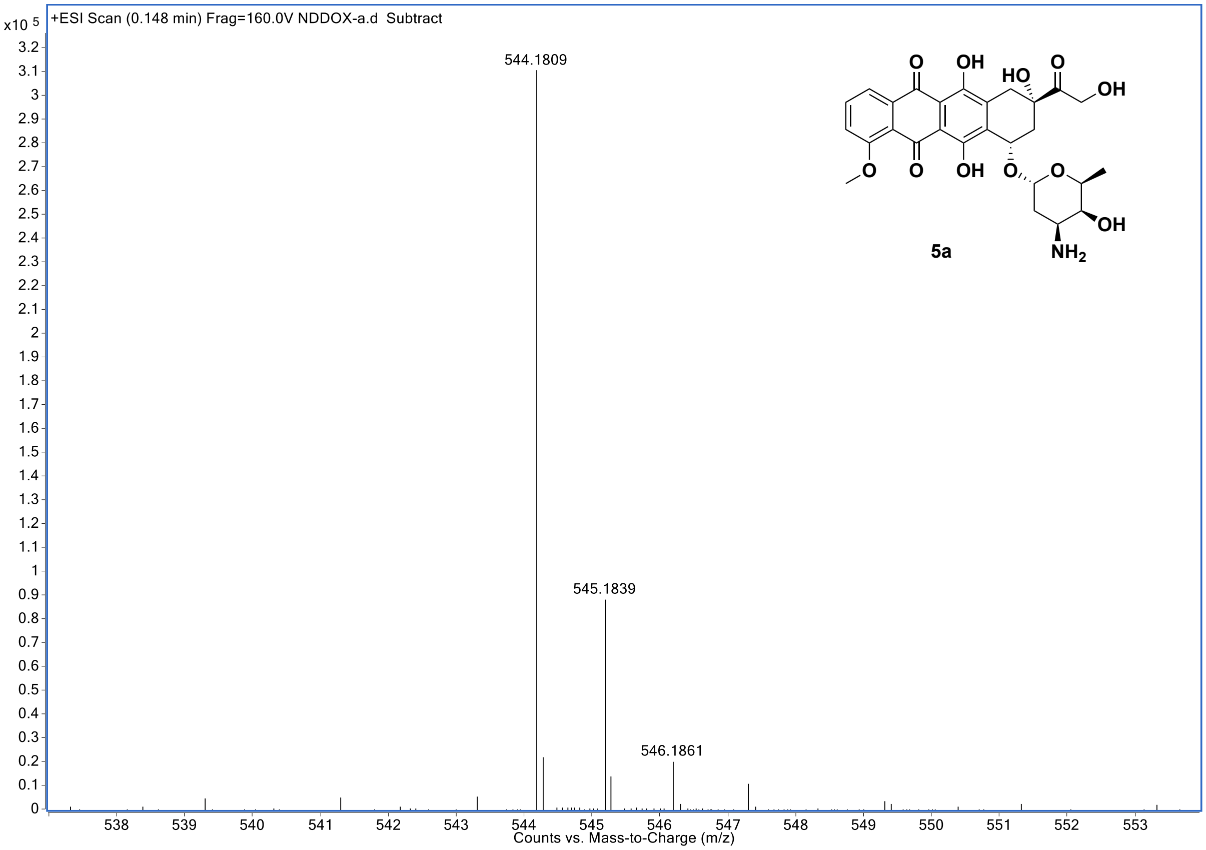

+ESI Scan (0.148 min) Frag=160.0V NDDOX-a.d Subtract
x10 5
544.1809
545.1839
546.1861
O
OH
HO
O
OH
O
O
OH
O
O
OH
NH2
5a
Counts vs. Mass-to-Charge (m/z)

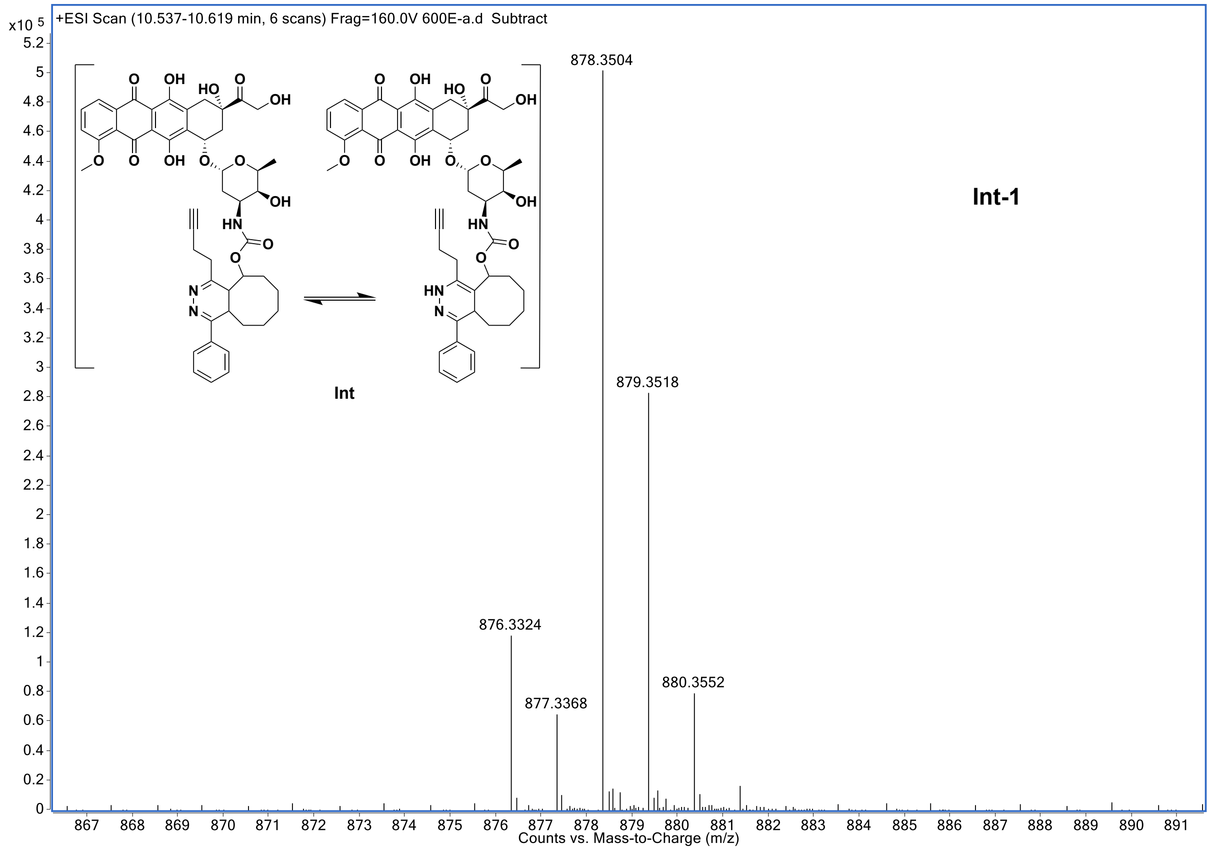

+ESI Scan (10.537-10.619 min, 6 scans) Frag=160.0V 600E-a.d Subtract
x10 5
878.3504
879.3518
876.3324
877.3368
880.3552
Int-1
Int
Counts vs. Mass-to-Charge (m/z)


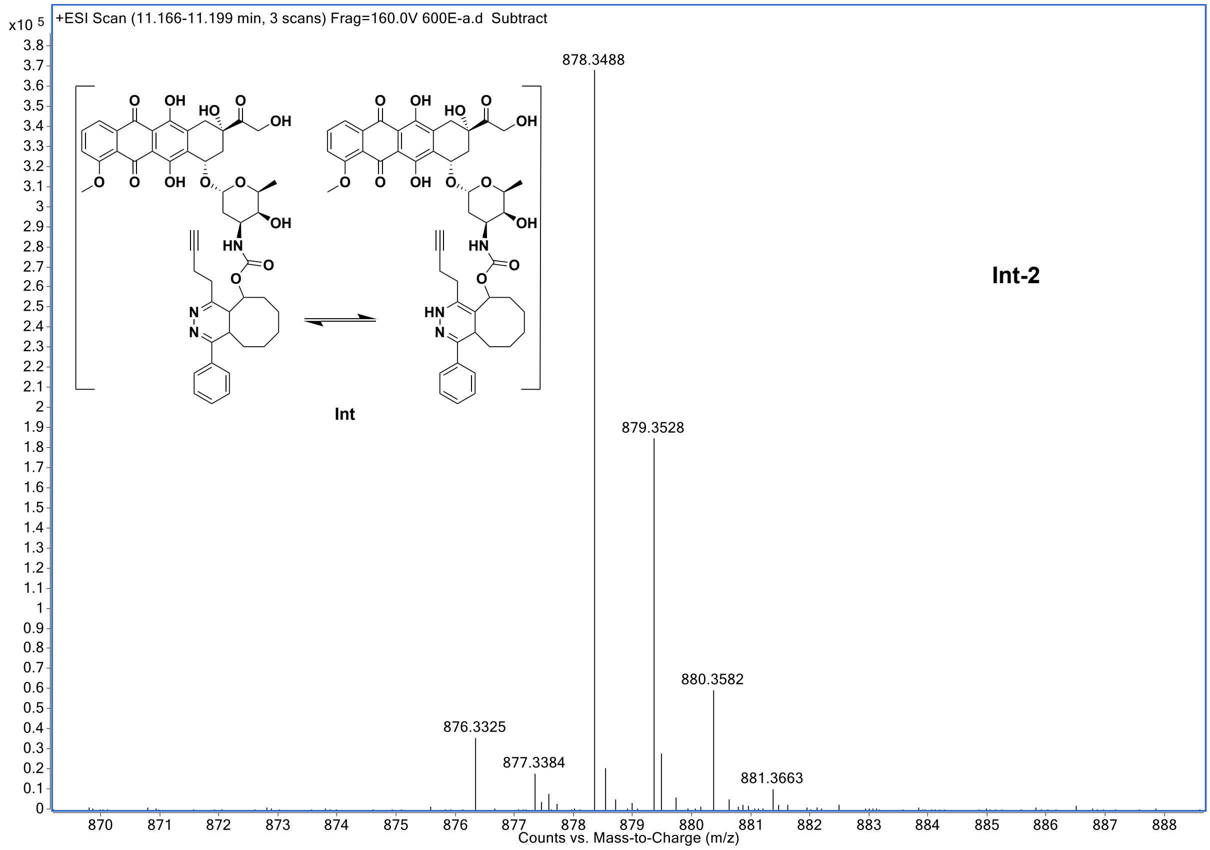

+ESI Scan (11.166-11.199 min, 3 scans) Frag=160.0V 600E-a.d Subtract
x10 5
878.3488
879.3528
880.3582
876.3325
877.3384
881.3663
Int-2
Int
Counts vs. Mass-to-Charge (m/z)

#### 7. The Stabilities of 1a and 2a under Fluorination Reaction Conditions

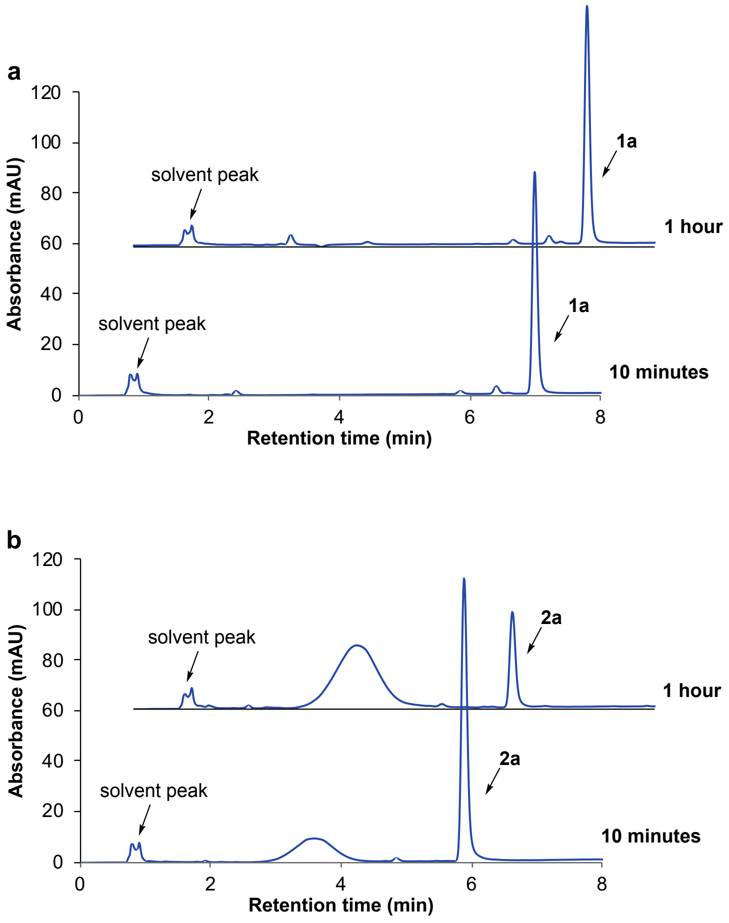


**Fig. 17. HPLC analysis of the stability of photocaged dihydrotetrazine 1a and tetrazine 2a under typical fluorination reaction conditions.** 100 µL of 3.5% aq. $K_2CO_3$ solution was added to a mixture of Kryptofix 222 (3.5 µmol) and KF (10 µmol) in MeCN (900 µL). Then the reaction mixture was heated to 85°C for 5 minutes. In the absence of light, photocaged dihydrotetrazine **1a** or tetrazine **2a** was added and the reaction mixture was heated for 1 hour at 85°C. **a,** A solution of photocaged dihydrotetrazine **1a** (3.2 µmol) in MeCN (1 mL) was added. **b,** A solution of tetrazine **2a** (3.2 µmol) in MeCN (1 mL) was added. Samples of the reaction mixture were taken at different time points (10 minutes and 1 hour) and examined by LC-MS.

HPLC analysis was carried out on an Eclipse Plus C8 analytical column with Phase A/Phase B gradients [Phase A: MeOH with 0.1% trifluoroacetic acid, Phase B: $H_2O$ with 0.1% trifluoroacetic acid]. 50% Phase A in Phase B, 1 minute, and 50%–70% Phase A in Phase B, 4 minutes, then 70%–90% Phase A in Phase B, 3 minutes. The absorbance was monitored at 280 nm.

As shown in Fig. 17, under typical fluorination reaction conditions, 38% of tetrazine **2a** degrades in 10 minutes and 79% of degradation was observed in 1 hour. While photocaged dihydrotetrazine **1a** is stable under the same reaction conditions and less than 5% of degradation was observed in 1 hour.

### 8. NMR Spectra

#### 1-(2-Nitrophenyl)ethyl 6-(but-3-yn-1-yl)-3-phenyl-1,2,4,5-tetrazine-1(4H)-carboxylate (1a)

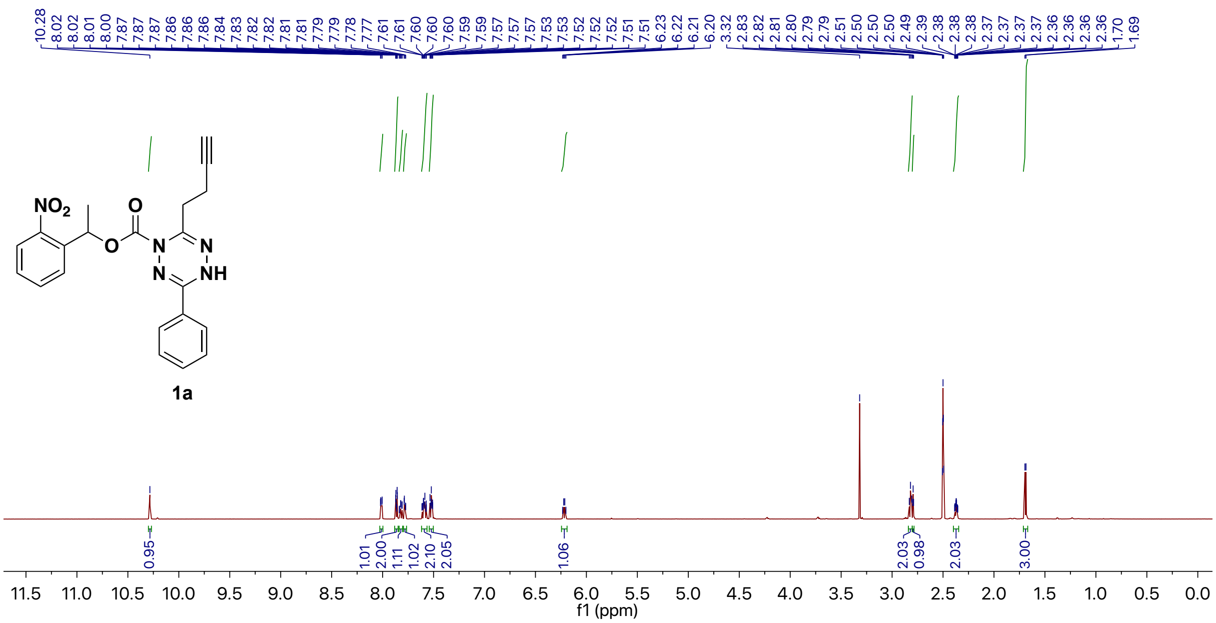


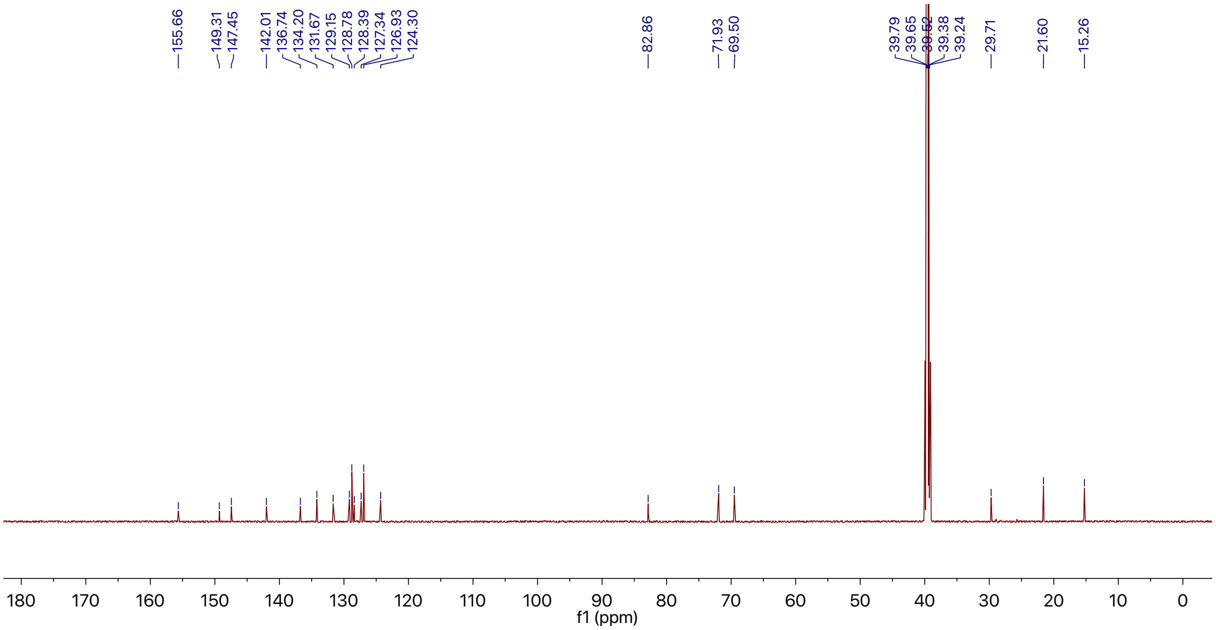

#### $^1$H-$^1$H-NEOSY NMR Spectrum

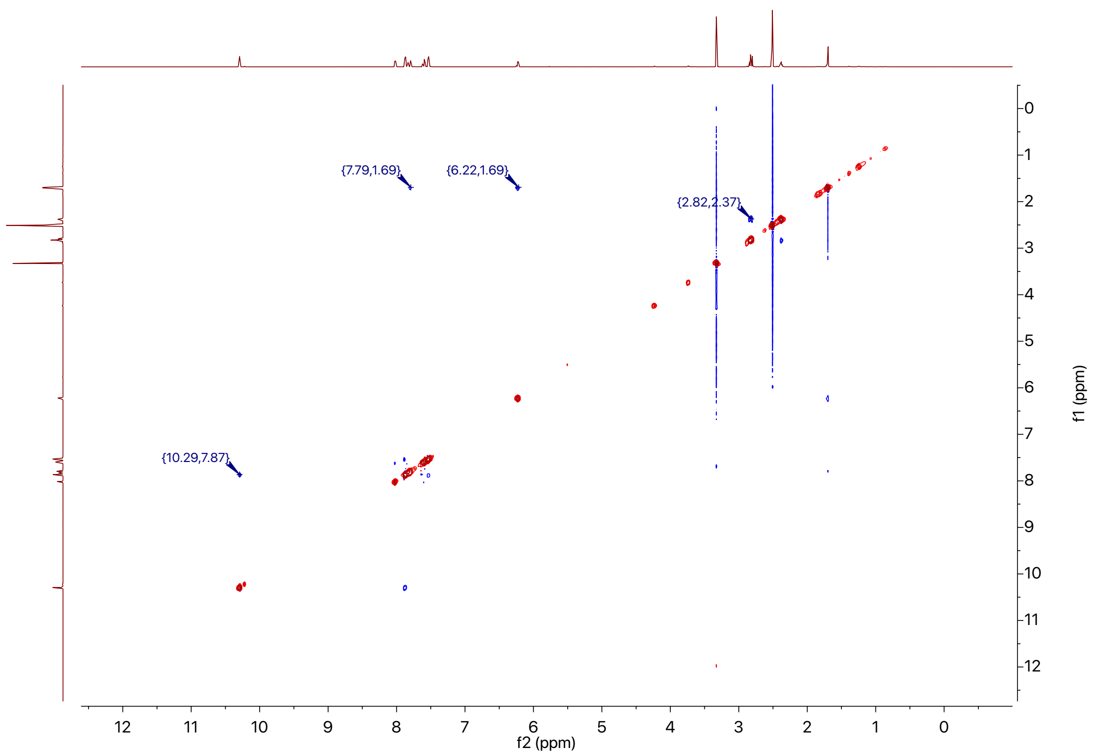


#### $^1$H-$^{13}$C-HSQC NMR Spectrum

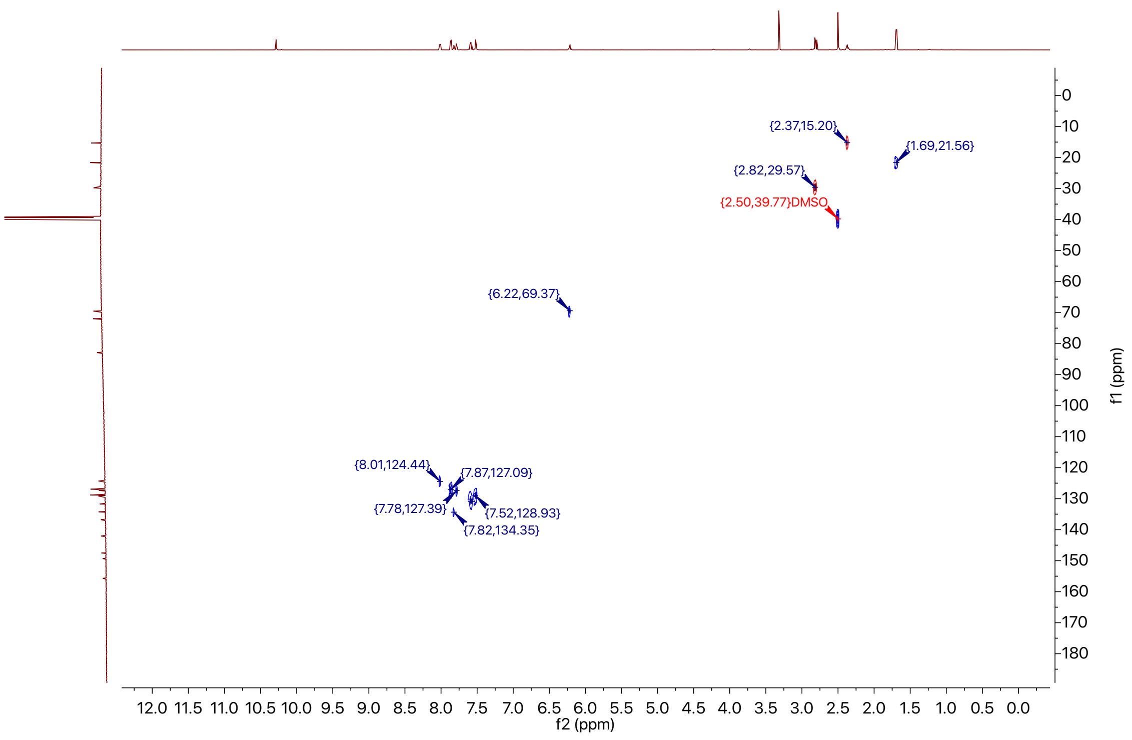

#### $^{1}H$-$^{15}N$ HSQC NMR Spectrum

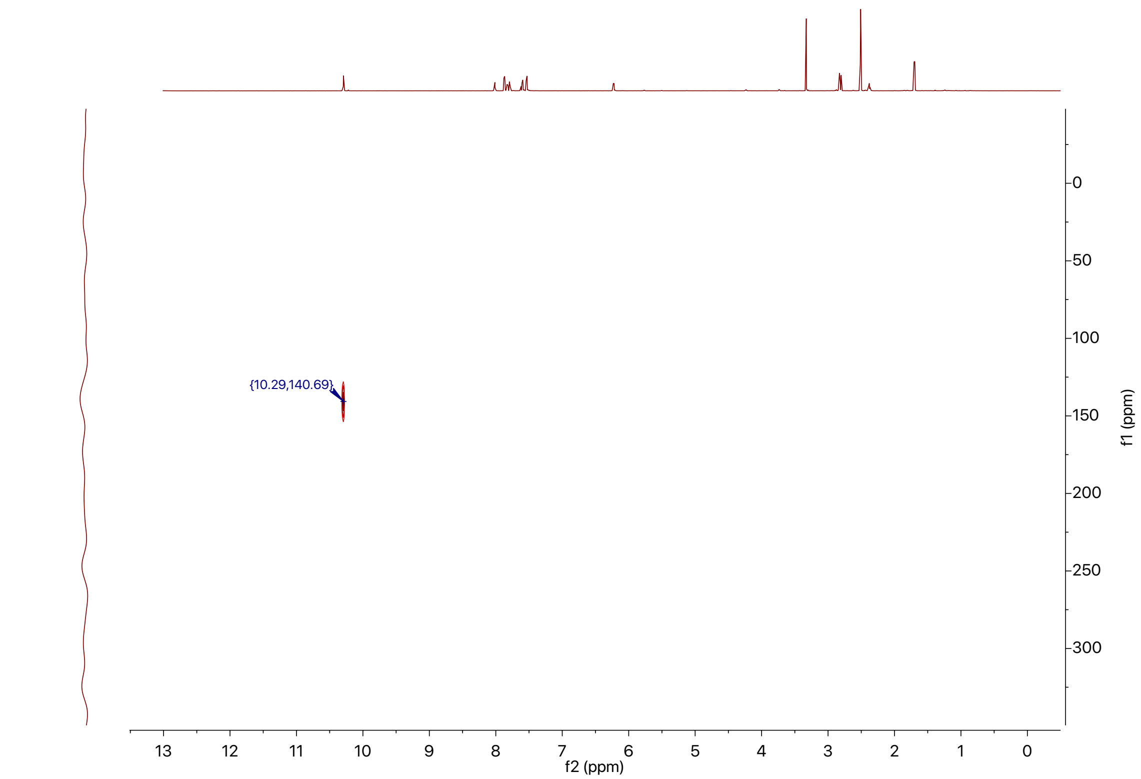


#### $^{1}H$-$^{13}C$-HMBC NMR Spectrum

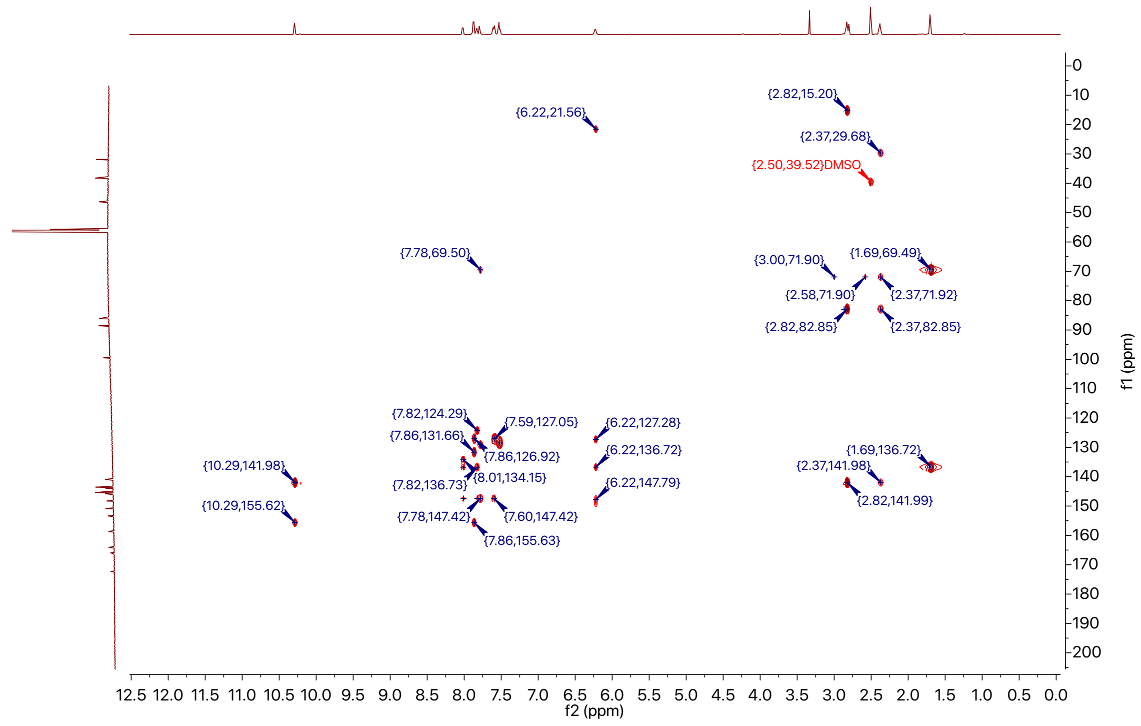

#### $^{1}H$-$^{5}N$-HMBC NMR Spectrum

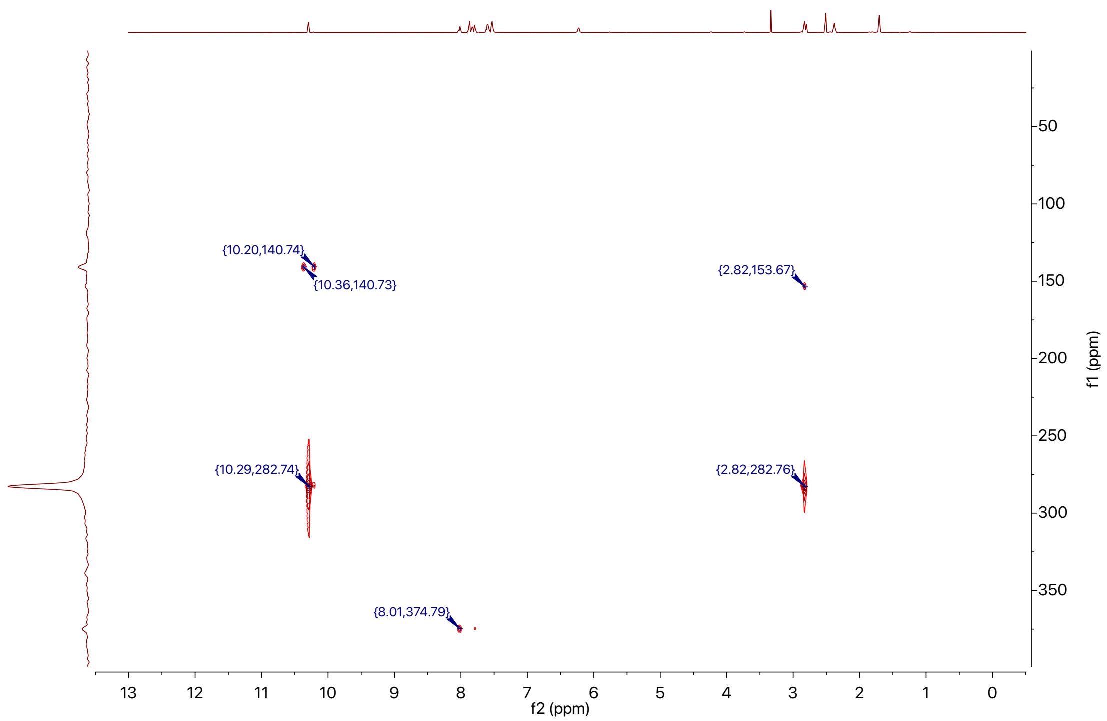

#### 3-(But-3-yn-1-yl)-6-phenyl-1,2,4,5-tetrazine (2a)

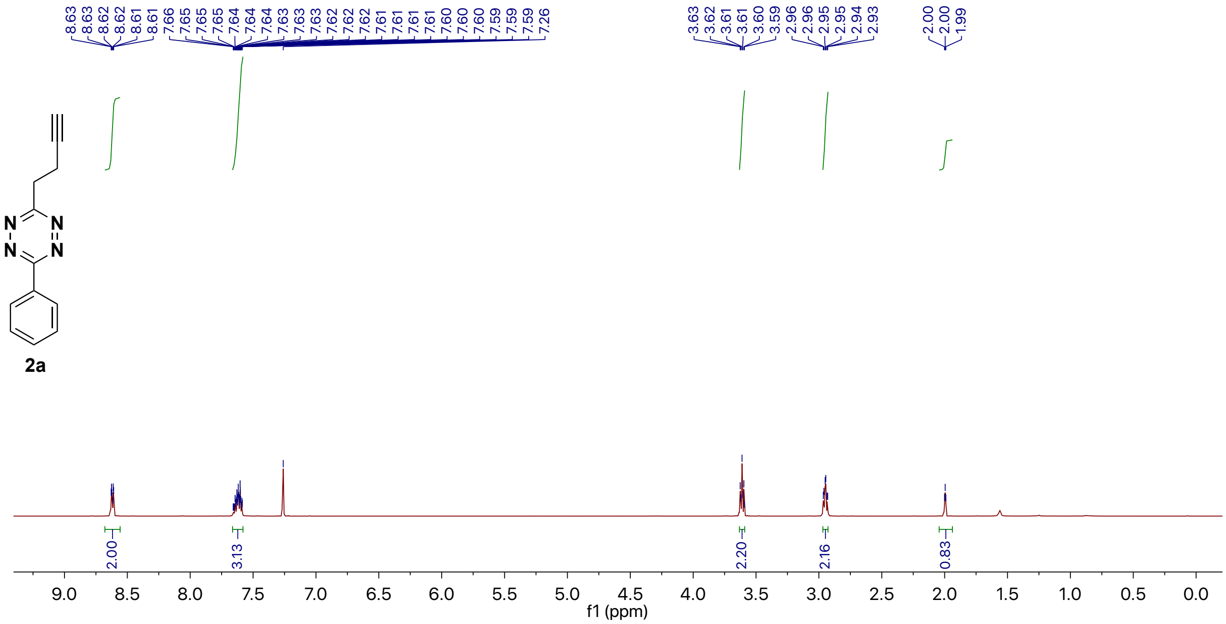


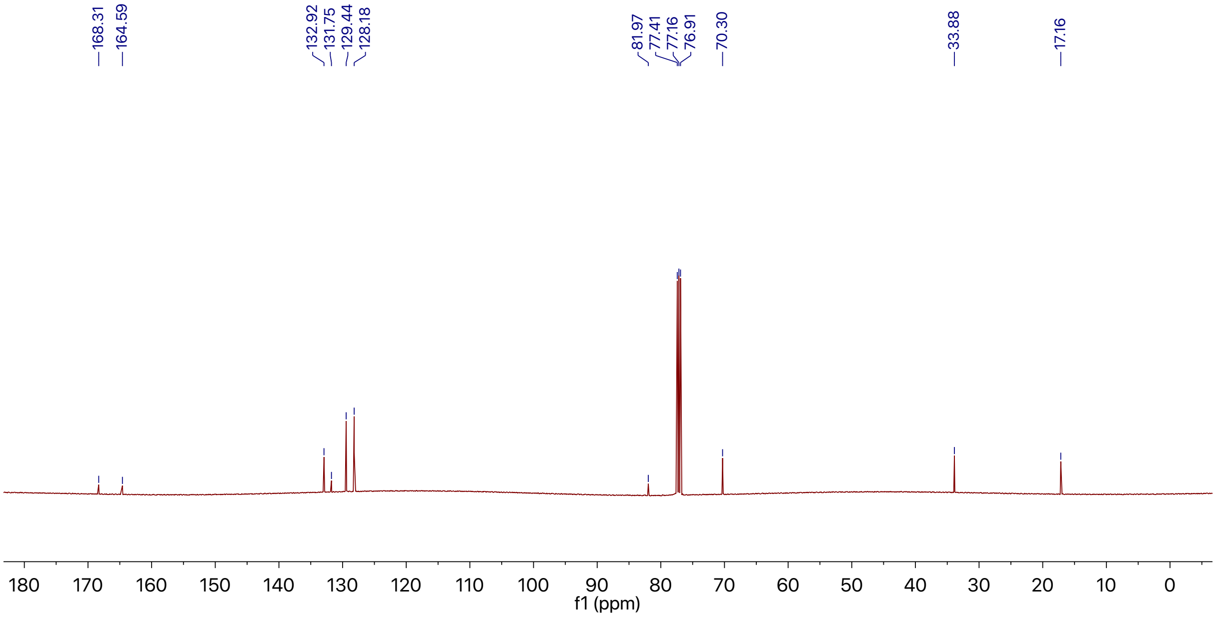

#### Dihydrotetrazine-Fmoc-lysine (1b)

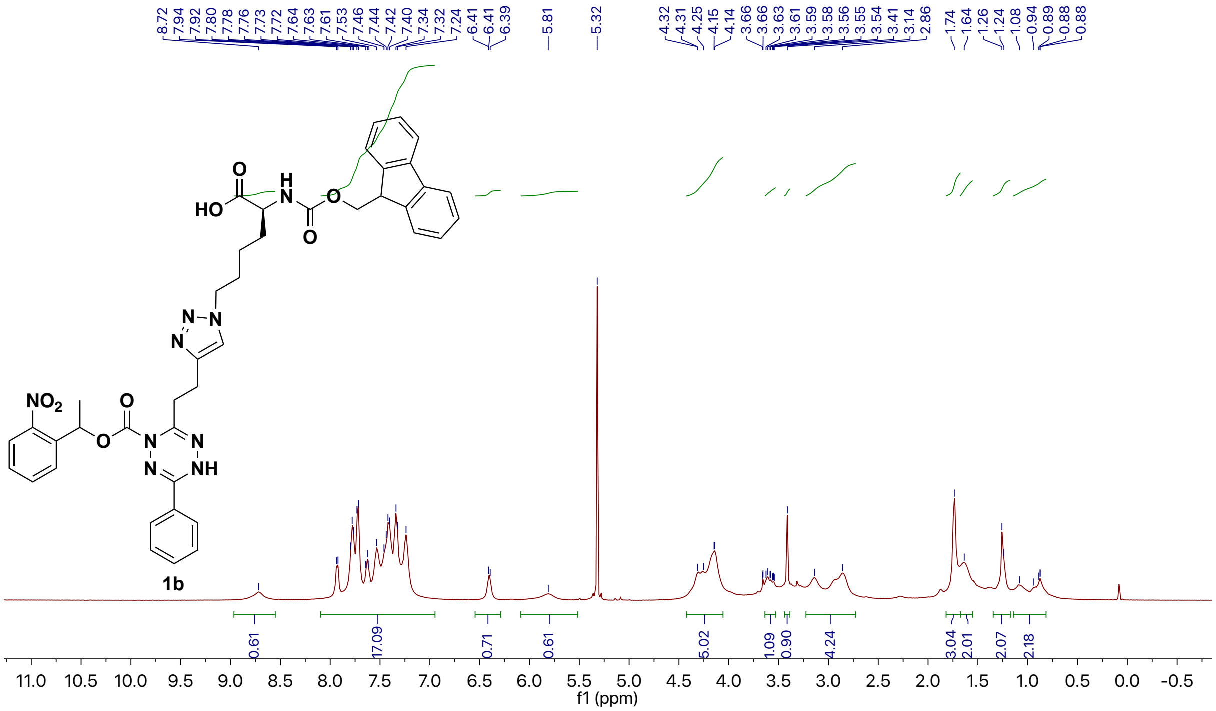


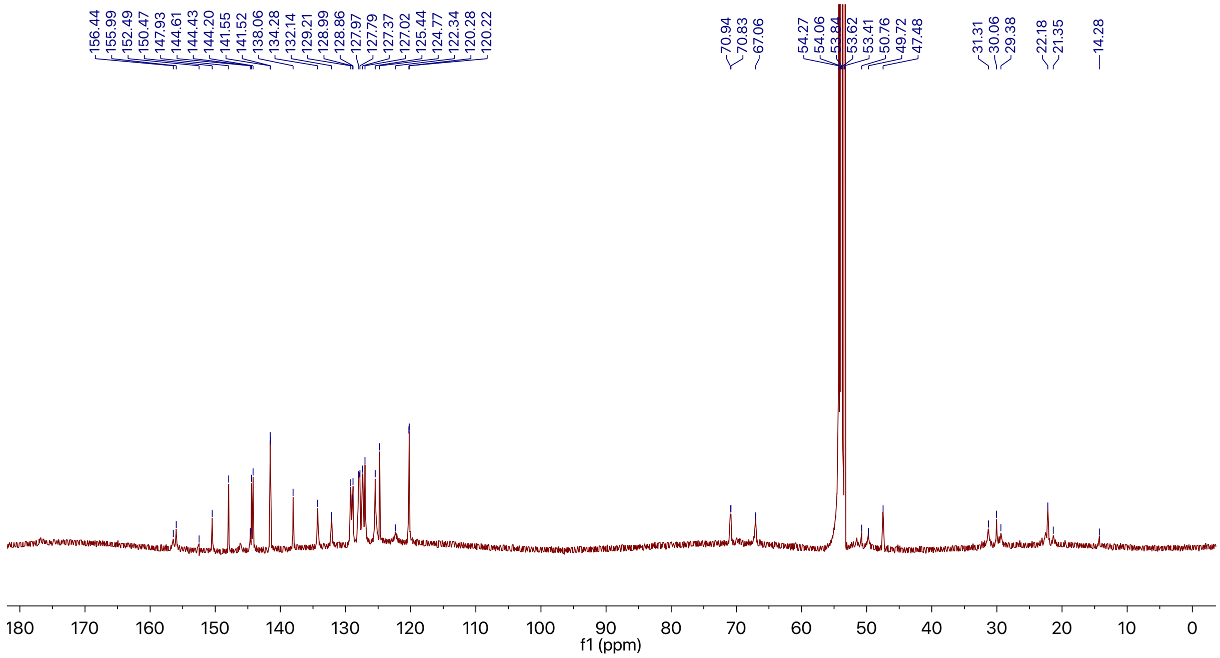

#### *Trans*-cyclooctene carbamate caged doxorubicin (3c)

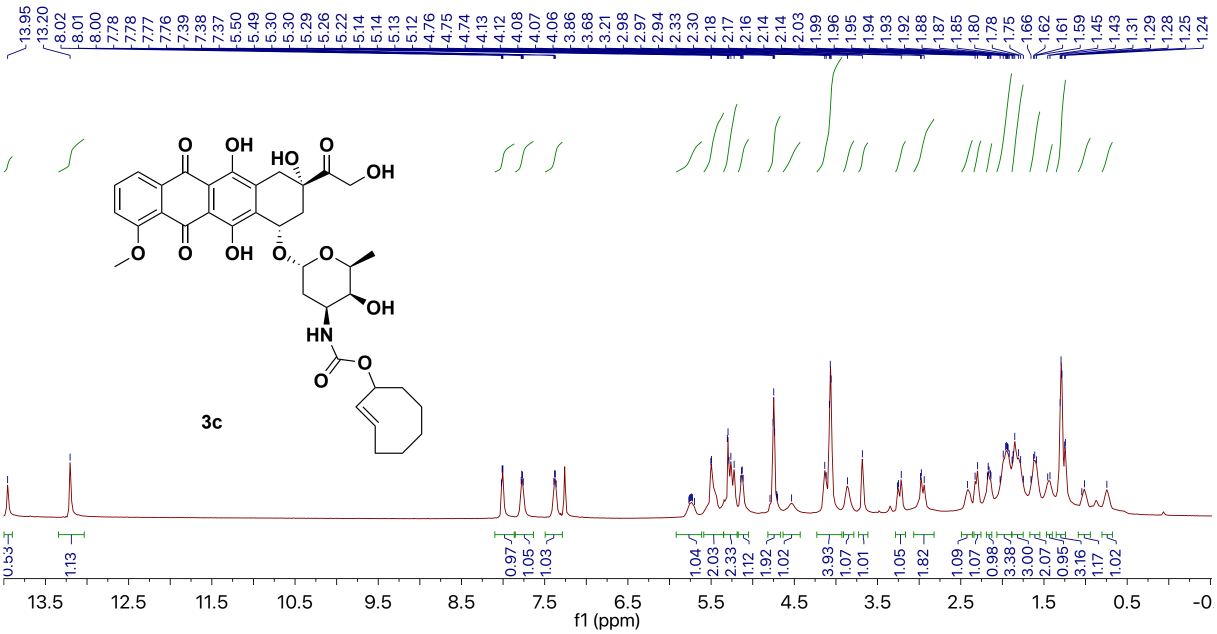


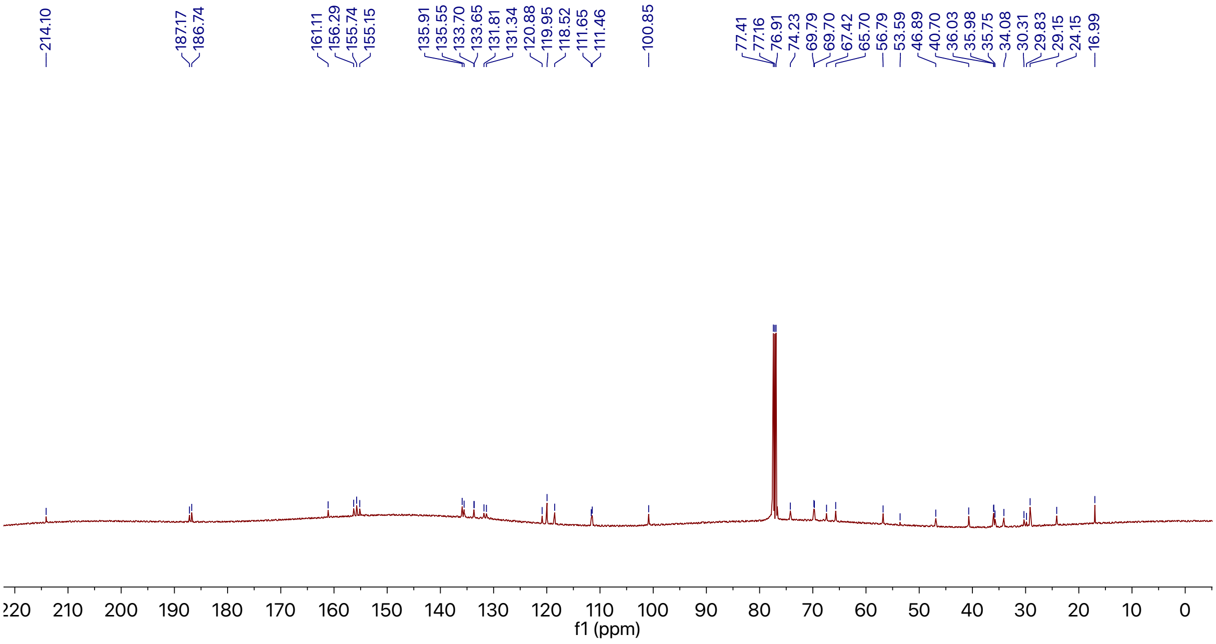

#### 9. References


1. Mao, W., Shi, W., Li, J., Su, D., Wang, X., Zhang, L., Pan, L., Wu, X. & Wu, H. Organocatalytic and scalable syntheses of unsymmetrical 1,2,4,5-tetrazines by thiol-containing promotors. *Angew. Chem. Int. Ed*. **58**, 1106–1109 (2019).

2. Makarov, S.V., Kudrik, E.V. & Davydov, K.A. Reaction of thiourea *S*,*S*-dioxides with dyes containing carbonyl or azo groups. *Russ. J. Gen. Chem.* **76**, 1599–1603 (2006).

3. Shiozaki, H., Miyahara, M., Otsuka, K., Miyako, K., Honda, A., Takasaki, Y., Takamizawa, S., Tukada, H., Ishikawa, Y., Sakai, R. & Oikawa, M. Studies on Aculeines: synthetic strategy to the fully protected Protoaculeine B, the *N*-terminal amino acid of Aculeine B. *Org. Lett.* **20**, 3403–3407 (2018).

4. Versteegen, R. M., ten Hoeve, W., Rossin, R., de Geus, M. A. R., Janssen, H. M. & Robillard, M. S. Click-to-release from *trans*-cyclooctenes: mechanistic insights and expansion of scope from established carbamate to remarkable ether cleavage. *Angew. Chem. Int. Ed*. **57**, 10494–10499 (2018).

5. Jin, S., Brea, R. J., Rudd, A. K., Moon, S. P., Pratt, M. R. & Devaraj, N. K. Traceless native chemical ligation of lipid-modified peptide surfactants by mixed micelle formation. *Nat. Commun.* **11**, 2793 (2020).

6. Eissler, S., Kley, M., Bächle, D., Loidl, G., Meier, T. & Samson, D. Substitution determination of Fmoc-substituted resins at different wavelengths. *J. Pept. Sci.* **23**, 757–762 (2017).
